# Thiol-ene cross-linked alginate hydrogel encapsulation modulates the extracellular matrix of kidney organoids by reducing abnormal type 1a1 collagen deposition

**DOI:** 10.1101/2020.11.17.386250

**Authors:** Thomas Geuens, Floor A.A. Ruiter, Anika Schumacher, Francis L. C. Morgan, Timo Rademakers, Loes E. Wiersma, Cathelijne W. van den Berg, Ton J. Rabelink, Matthew B. Baker, Vanessa L.S. LaPointe

**Author notes:** Equally contributed (Thomas Geuens, Floor A.A. Ruiter). Corresponding authors To whom correspondence should be addressed: Dr. Vanessa LS LaPointe and Dr. Matthew B. Baker, MERLN Institute for Technology-Inspired Regenerative Medicine, Maastricht University P.O. Box 616, 6200 MD, Maastricht, the Netherlands, and. **CRediT author statement Thomas Geuens:** Conceptualization, Performing of experiments, Investigation, Data analyses, Writing. **Floor A.A. Ruiter:** Conceptualization, Performing of experiments, Investigation, Data analyses, Writing. **Anika Schumacher:** Performing of experiments. **Francis L. C. Morgan:** Performing of experiments, Data analyses. **Timo Rademakers:** Performing of experiments. **Loes E. Wiersma:** Experimental Support. **Cathelijne W. van den Berg:** Experimental Support, Supervision. **Ton J. Rabelink:** Experimental Support, Supervision. **Matthew B. Baker:** Conceptualization, Reviewing, Editing, Supervision. **Vanessa L.S. LaPointe:** Conceptualization, Writing, Reviewing, Editing, Supervision.

## Abstract

Differentiated kidney organoids from induced pluripotent stem cells hold promise as a treatment for patients with kidney diseases. Before these organoids can be translated to the clinic, shortcomings regarding their cellular, extracellular compositions and developmental plateau needs to be overcome. We performed a proteomic analysis on kidney organoids cultured for a prolonged culture time and we found a specific change in the extracellular matrix composition with increased expression of types 1a1, 2 and 6a1 collagen. Such an excessive accumulation of specific collagen types is a hallmark of renal fibrosis that causes a life-threatening pathological condition by compromising key functions of the human kidney. Here we hypothesized the need for a three-dimensional environment to grow the kidney organoids, which could better mimic the in vivo surroundings of the developing kidney than standard culture on a transwell filter. Encapsulating organoids for four days in a soft, thiol-ene cross-linked alginate hydrogel resulted in decreased type 1a1 collagen expression. Furthermore, the encapsulation did not result in any changes of organoid structural morphology. Using a biomaterial to modulate collagen expression allows for a prolonged kidney organoid culture in vitro and a reduction of abnormal type 1a1 collagen expression bringing kidney organoids closer to clinical application.

**HIGHLIGHTS:**

- Prolonging kidney organoid culture results in a developmental plateau instead of improved *in vitro* maturation.
- Proteomic analyses point to an increased expression of specific collagen subtypes associated with renal fibrosis.
- Encapsulating kidney organoids using a soft thiol-ene cross-linked alginate hydrogel reduces collagen type 1a1 and αSMA deposition.

## INTRODUCTION

With chronic kidney disease (CKD) reaching epidemic proportions (affecting 11–13% of the worldwide population), there is an increasing need for regenerative strategies for end-stage renal diseases [1]. There are currently two million people depending on dialysis with no option for renal transplantation [2], and the few who are eligible for kidney transplants face a high chance of rejection, therefore further diminishing the number of successful treatments [2, 3].

One regenerative medicine-based approach is to provide an alternative solution in the form of a kidney organoid developed in vitro that could serve as a functional engraftment to the dysfunctional kidney [4, 5]. For example, by using patient-derived human induced pluripotent stem cells (hiPSCs), patient-specific solutions can be generated. Using knowledge of development, hiPSCs have been directed into organoid cultures that are being employed as model systems for various human organs [6], including the mimic of kidney organogenesis to generate renal organoids [7–11]. Despite these major breakthroughs, there are still challenges with the current state-of-the-art and their therapeutic suitability [12]. For instance, while the kidney organoids recapitulate many of the structural elements observed in a developing human kidney, they still face immaturity at both a transcriptional [13] and morphological level [14].

Here we show that prolonging kidney organoid culture in vitro does not improve organoid maturity but instead negatively impacts the extracellular matrix (ECM) formation, characterized by the increase of specific types of collagens, such as types 1a1, 2 and 6a1, which are hallmarks of fibrosis. Such extracellular modifications have been associated with pathological conditions ultimately compromising key renal functions, such as blood filtration and urine production [15–17]. We hypothesized the need for a three-dimensional (3D) hydrogel environment to more closely mimic the in vivo ECM. Therefore, we designed a thiol-ene cross-linked alginate hydrogel to encapsulate kidney organoids during part of the culture period, which resulted in a reduced deposition of type 1a1 collagen with no changes to the renal structures. Including this hydrogel as part of the in vitro culture protocol can therefore improve the phenotype of kidney organoids, making them more suitable for regenerative medicine applications.

## MATERIALS AND METHODS

### hiPSC culture

hiPSCs were expanded on vitronectin-coated (0.5 μg/cm2) plates in E8 medium (Thermo Fisher Scientific) and were passaged twice weekly using TrypLE Express (Thermo Fisher Scientific) with the addition of RevitaCell Supplement (Thermo Fisher Scientific) for 24h. The hiPSC line LUMC0072iCTRL01 was obtained from the hiPSC core facility at the Leiden University Medical Center where they were generated from fibroblasts using the Simplicon RNA reprogramming kit (Millipore).

### Differentiation and organoid formation

Kidney organoids were generated from hiPSCs according to an established protocol (10). Briefly, hiPSCs were seeded on vitronectin-coated (0.5 μg/cm2) plates at 7,300 cells/cm2 in E8 medium supplemented with RevitaCell Supplement. After 24 h, differentiation was started (day 0) by changing to STEMdiff APEL medium (STEMCELL Technologies) supplemented with 8 μM CHIR99021 (R&D Systems), 1% v/v antibiotic/antimycotic (Thermo Fisher Scientific), and 1% (v/v) PFHM-II protein-free hybridoma medium (Thermo Fisher Scientific). On day 4 of differentiation, medium was switched to STEMdiff APEL medium supplemented with 200 ng/μl FGF-9 (R&D Systems) and 1 μg/ml heparin (Sigma-Aldrich). By day 7, the cells formed a confluent monolayer and were subjected to a 1 h pulse with 5 μM CHIR99021 before being harvested by trypsinization. An aggregate of 500,000 cells was transferred to a transwell filter with a 0.4 μm pore size (Corning). From this moment onwards, kidney organoids were cultured for 4 more days (termed day 7+4) in STEMdiff APEL2 medium supplemented with FGF-9 and heparin. Beginning on day 7+5, STEMdiff APEL2 medium without supplemented growth factors was changed every 2 days. The organoids were maintained on the filter membranes until day 7+18 or day 7+25.

### Organoid decellularization

All incubation and centrifugation steps were performed at 4°C to minimize proteolysis. Kidney organoids were harvested from the filter membrane and snap frozen in liquid nitrogen for storage or used immediately for decellularization. Organoids were incubated for 30 min in an extraction buffer containing Tris-HCl (pH 7.8; 10 mM), NaCl (150 mM), EDTA (25 mM), 1% (v/v) Triton X-100 and one tablet of protease inhibitors (Roche). Samples were centrifuged at 14,000 × g for 10 min to yield the supernatant termed fraction one. The remaining pellet was incubated for 30 min in an alkaline detergent buffer containing NH4OH (20 mM) and 0.5% (v/v) Triton X-100 diluted in PBS. Samples were centrifuged at 14,000 × g for 10 min to yield fraction two. The remaining pellet was incubated for 30 min in DNase I buffer containing 25 units DNase I diluted in 50 μl PBS. Samples were centrifuged at 14,000 × g for 10 min to yield fraction three. The final remaining pellet was resuspended in Laemmli sample buffer (Bio-Rad) with 1,4-dithiothreitol (DTT; 100 mM). Before the samples were subjected to heat denaturation at 95°C for 10 min, they were sonicated three times for 5 sec with 10% amplitude and 30 sec between each sonication round.

### Mass spectrometry

Protein samples were resolved by SDS-PAGE (4–12%) and visualized by Coomassie staining (0.05% (w/v) Coomassie R-250). Samples were run onto the interface of the stacking and running gel, manually excised from the gel and subjected to in-gel digestion using a MassPREP digestion robot (Waters, Manchester UK). Briefly, a solution of NH_4_HCO_3_ (50 mM) in 50% (v/v) acetonitrile (ACN) was used for destaining the gel slices. Cysteines were reduced using DTT (10 mM) in NH_4_HCO_3_ (100 mM) for 30 min followed by an alkylation reaction using iodoacetamide (55 mM) in NH_4_HCO_3_ (100 mM) for 20 min. Afterwards, samples were washed with NH_4_HCO_3_ (100 mM) and subsequently dehydrated with 100% (v/v) ACN. Trypsin (6 ng/ml, Promega) in NH_4_HCO_3_ (50 mM) was added and incubated at 37°C for 5 h. Peptides were extracted using 1% (v/v) formic acid (FA) supplemented with 2% (v/v) ACN. A subsequent extraction step was performed using 1% (v/v) FA supplemented with 50% (v/v) ACN.

Once extracted, peptides were separated on an Ultimate 3000 Rapid Separation UHPLC (Thermo Fisher Scientific) equipped with an C18 analytical column (2 μm particle size, 100 Å pore size, 75 μm inner diameter, 150 mm length; Acclaim). Peptide samples were desalted and separated on the C18 analytical column with a 90 min linear gradient from 5– 35% (v/v) ACN with 0.1% (v/v) FA at a 300 nl/min flow rate. This column was coupled online to a Q Exactive HF mass spectrometer (Thermo Fisher Scientific) using a full MS scan from 350–1,650 mz-1 at a resolution of 120,000. Subsequent MS/MS scans of the top 15 most intense ions were done at a resolution of 15,000.

### Mass spectrometry data analyses

The obtained spectra were analyzed using the Proteome Discoverer software (version 2.2, Thermo Fisher Scientific) using the Sequest search engine and the SwissProt Human database (TaxID=9606, v2017-10-25). The database search was performed with the following settings: trypsin digestion with a maximum of two missed cleavages, minimum peptide length of six amino acids, precursor mass tolerance of 10 ppm, fragment mass tolerance of 0.02 Da, dynamic modifications of methionine oxidation and protein N-terminus acetylation, static modification of cysteine carbamidomethylation. Data analyses were performed using the Perseus software platform [18].

### Western blot

Protein samples were resolved by SDS-PAGE (Bio-Rad) and transferred to a PVDF membrane (Bio-Rad). Blocking was performed using 5% (w/v) milk powder diluted in TBS supplemented with 0.1% (v/v) Tween-20 for 1 h. Membranes were incubated with primary antibodies overnight at 4°C and for 1 h at room temperature (RT) with secondary horseradish peroxidase conjugated antibodies (Bio-Rad, 1:3000). Membranes were then developed using the Clarity Western ECL Substrate (Bio-Rad) and imaged on a ChemiDoc MP Imaging System (Bio-Rad). The following antibodies were used: type 1a1 collagen (ab6308, Abcam, 1:1000), type 6a1 collagen (GTX109963, Genetex, 1:3000), laminin B2 (sc-59980, Santa Cruz Biotechnology, 1:500), nidogen (sc-133175, Santa Cruz Biotechnology, 1:500), beta-actin (ab8227 Abcam, 1:3000) and GAPDH (11335232, Thermo Fisher Scientific, 1:3000).

### Analysis of off-target gene expression in iPSC-derived kidney organoids

Previously generated single-cell RNAseq data of iPSC-derived kidney organoids using the Takasato protocol (13) were downloaded from the Gene Expression Omnibus (GEO: GSE118184). The raw transcript count tables were analyzed separately for each time point (day 0, 7, 7+5, 7+12, 7+19 and 7+27) using R software (3.6.2) and the Seurat package (version 3.2.0). Low quality cells were previously excluded, as described by Wu et al. (13). The gene expression matrices were normalized for sequencing depth per cell and log-transformed using a scaling factor of 10,000. Then 2,000 features with the highest cell-to-cell variation were identified, scaled and centered and used for principal component analysis. The principal components representing the true dimensionality of each dataset were selected and further used for non-linear dimensional reduction. Normalized gene expression of markers of interest are shown in tSNE space.

### Norbornene-alginate (NB-Alg) synthesis

Purified sodium alginate (0.5 g, 2.5 mmol COOH groups, 1 equiv., Manugel GMB, FMC, lot no. G940200) was dissolved overnight in 50 ml MES buffer (0.1 M MES, 0.3 M NaCl, pH 6.5) as previously described (19). 1-(3-(Dimethylamino)propyl)-3-ethylcarboiimide hydrochloride (EDC-HCL; 0.5 g, 2.6 mmol, 1.02 equiv., VWR) and NHS-ester (0.65 g, 5.65 mmol, 2.3 equiv., Sigma-Aldrich) were added and the solution was stirred for 30 min. The pH of the solution was adjusted to approximately 8 (with 5 M HCl) before the addition of the 5-norbornene-2-methylamine (9.4 × 10-2 g, 0.76 mmol, 0.3 equiv., mixture of isomers, TCI Chemicals) and was stirred overnight (18 h) at RT. The norbornene-functionalized alginate product was purified by dialysis in a 10 kDa MWCO dialysis tube (Spectra/Por, regenerated cellulose, VWR) in 100 mM, 50 mM, 25 mM and 0 mM NaCl in MilliQ water, which was changed every 10–18 h (dialysis ratio 1:50). The resultant product was freeze-dried and its structure confirmed by H1-NMR in deuterated water (D2O, Figure S5). The percentage of norbornene functionalization (3.5%) was calculated relative to the DMF internal standard in the H1-NMR spectra (Table S1). MW of the NB-Alg was determined by GPC (Figure S6b-c and Table S2)

### ^1^H-NMR analysis

^1^H-NMR spectra were recorded on a Bruker AVANCE III HD 700-MHz spectrometer equipped with a cryogenically cooled three-channel TCI probe in D_2_O with dimethylformaldehyde (DMF) as an internal standard (0.2 M). A water suppression pulse sequence was applied to spectra. MestReNova 11.0 software was used to analyse the obtained spectra. All chemical shifts are reported in parts per million (ppm) relative to DMF (H(C=O), 8 ppm).

### GPC analysis

Molecular weight were measure in 0.1 M NaNO_3_ H_2_O eluent. Samples were eluted at RT at a flow rate of 0.5 mL/min on a Prominence-I LC-2030C3D LC (Shimadzu Europa GmbH) and Shodex SB-803/SB-804 HQ columns (Showax Denko America, Inc). Calibration was performed using PEG standards of molecular weights up to 300,000 MW (Agilent PEG calibration kit, Agilent Technologies). Molecular weight and dispersity values were calculated using LabSolutions GPC software (Shimadzu Europa GmbH). The samples were dissolved in 0.1 M NaNO_3_ H_2_O in a concentration of 0.5 mg/mL and filtered through 0.45 μm filters to remove any unwanted impurities.

### Organoid hydrogel encapsulation

Norbornene functionalized alginate (71 mg, 3.5% functionalization, 11.3 μmol norbornene units) was dissolved in 2.3 ml STEMdiff APEL2 medium. The 4-arm 10 kDa PEG thiol (5 mg, 0.02 mmol SH units, Creative PEGWorks) and LAP UV initiator (3.3 mg, 11.2 μMol, 3.2 mM, Sigma-Aldrich) were dissolved separately in 1.2 ml APEL2 medium. Both solutions were passed through a 0.2 μm sterilization filter. The cross-linking/initiator solution was added to the norbornene alginate solution and stirred for 30 min before organoid encapsulation to form a 2% (w/v) NB-alginate solution. The solution was deposited over the organoids on the top of the transwell membrane at day 7+14 or day 7+21 of culture and exposed to 365 nm light (10 mW/cm2, 30 sec, UVP CL-1000 ultraviolet cross-linker). After cross-linking, APEL2 medium (1.2 ml) was added to the bottom of the transwell filters, and the organoids were cultured for three additional days until day 7+18 or 7+25.

### Rheometry

Rheometry was performed on a TA instruments DHR-2 rheometer with a 20 mm cone-plate geometry and a cone angle of 2.002°. The hydrogel for these experiments did not contain organoids. Precursor solutions were equilibrated at RT, and the temperature was maintained at 23–24°C. Time sweeps at 1% strain and 10 rad/s were performed for 10 min total. After 5 min of measurement to ensure stability, the sample was irradiated with 365 nm UV light at 10 mW/cm2 for 2 min. The time sweep was prolonged for another 3 min to ensure no short-term evolution following cross-linking. The same sample was then subjected to a frequency sweep from 100–0.1 rad/s at 1% strain, followed by a strain sweep from 1–1000% at 10 rad/s. Each measurement was performed in triplicate.

### Swelling test

A swelling test was performed on transwell filters. The hydrogel solution was prepared and added on the transwell without organoids. APEL2 medium was added under the transwell filter and incubated for 98 h at 37°C. The weight of the transwell filter with the hydrogel was measured at 1, 2, 4, 6, 24, 48, 72 and 96 h. The swelling ratio in % (Sr) over time was calculated by the following equation: S_r=(w_1-w_o)/w_o, in which, w0 is the initial weight of the hydrogel and w1 is the weight of hydrogel at the specific time point.

### Cell viability assay

Cell viability was analyzed by EthD1/calcein AM staining. The medium was aspirated and the encapsulating hydrogel was removed and replaced with EthD1 (4 μM) and calcein AM (2 μM) in PBS and incubated for 30 min at RT. For cryo-sectioning, organoids were subsequently fixed with 2% (v/v) PFA for 10 min at 4°C and stored in PBS.

### Immunohistochemistry

Organoids were fixed in 4% (w/v) paraformaldehyde (PFA) for 20 min at 4°C, after which they were dehydrated by an overnight incubation in PBS containing 15% (w/v) sucrose followed by a second overnight incubation with 30% (w/v) sucrose. The next day, organoids were embedded in freezing solution containing 15% (w/v) sucrose and 7.5% (w/v) gelatin in PBS. The cast was placed in a beaker with isopentane and left to freeze for several minutes in liquid N2. Frozen organoids were stored at −20°C until cryo-sectioning, for which they were sectioned to 20 μm thickness at −18°C. Frozen organoid sections were placed in Coplin jars containing PBS and warmed to 37°C for 15–20 min to remove the embedding solution. Once removed, they were washed in PBS and permeabilized with 0.5% v/v IGEPAL in PBS for 15 min at RT and were blocked with 5% w/v BSA in PBS for 20 min at RT. The slides were then incubated overnight in the dark at 4°C with primary antibodies against: Nephrin (NPHS1,AF4269, R&D Systems, 1:300), Lotus Tetragonolobud Lectin (LTL, B-1325, Vectorlabs, 1:300), MEIS1/2/3 (sc-101850, Santa Cruz Biotechnology, 1:300), E-cadherin (ECAD, 610181, BD Biosciences, 1:300), SLC12A1 (HPA01496, Thermo Fisher Scientific, 1:200), collagen I (ab6308, Abcam, 1:300), type 6a1 collagen (GTX109963, Genetex, 1:300), α-Smooth muscle actin (aSMA, A2547, Sigma Aldrich, 1:100) and Fibronectin (FN, NBP1-91258, Novus Biologicals, 1:400). Subsequently, slides were washed and incubated with goat anti rabbit and goat anti mouse secondary antibodies for 45 min at RT in the dark: Alexa Fluor 488, Alexa Fluor 568 and Streptavidin Alexa Fluor 647 (Thermo Fisher Scientific, 1:100). After washing, nuclei were counterstained with DAPI (0.1 μg/ml) for 5 min.

For trichrome staining, organoids were fixed in 4% PFA for 20 min at 4°C and embedded in paraffin. Afterwards, 4 μm sections were cut and deparaffinized in xylene followed by a series of hydration steps. Trichrome staining was then performed using the Trichrome Stain (Masson) Kit (HT15, Sigma-Aldrich) following the manufacturer’s protocol. Mounted slides were imaged with an automated Nikon Eclipse Ti2-E microscope using a 10×, 20× or 40× air objective.

### Immunocytochemistry

hiPSCs were seeded on vitronectin-coated (0.5 μg/cm2) glass coverslips in a 24-well plate. Once sub-confluent, the cells were fixed with 4% (v/v) PFA for 20 min at RT. After three washes with PBS, they were permeabilized with 0.5% (v/v) Triton X-100 in PBS. The samples were then blocked for 1 h with 3% (v/v) goat serum in PBS. Subsequently, the samples were washed with 0.05% (v/v) Triton X-100 in PBS and incubated overnight at 4°C in the dark with the following primary antibodies against: OCT3/4 (sc-5279, Santa Cruz Biotechnology, 1:100), SOX2 (sc-365823, Santa Cruz Biotechnology, 1:100), SSEA4 (sc-21704, Santa Cruz Biotechnology, 1:100) and TRA-1-60 (sc-21705, Santa Cruz Biotechnology, 1:100). After overnight incubation, samples were washed three times with PBS and incubated with goat anti rabbit and goat anti mouse secondary antibodies for (Alexa Fluor 568) 1 h with (Thermo Fisher Scientific, 1:100). After washing, the nuclei were counterstained with DAPI (0.1 μg/ml) for 5 min. Mounted slides were imaged with an automated Nikon Eclipse Ti2-E microscope using a 20× or 40× air objective.

### Flow cytometry

hiPSCs were harvested using TrypLE Express (Thermo Fisher Scientific). One million cells were added to each tube, centrifuged for 4 min at 200 × g and resuspended in 100 μl PBS. The cells were fixed using Medium A from the FIX & PERM Cell Permeabilization Kit (Thermo Fisher Scientific) and incubated for 15 min at RT. They were washed with wash buffer (0.1% (v/v) sodium azide and 3% (w/v) BSA in PBS) and were then incubated with the primary antibodies for 20 min at RT. Those for extracellular antigens were diluted in wash buffer: SSEA4 (sc-21704, Santa Cruz Biotechnology, 1:100) and TRA-1-60 (sc-21705, Santa Cruz Biotechnology, 1:100). Those for intracellular antigens were diluted in Medium B for permeabilization: OCT4 (ab19857, Abcam, 1:100) and SOX2 (sc-365823, Santa Cruz Biotechnology, 1:100). An Alexa Fluor 488 (Thermo Fisher Scientific, 1:100) secondary antibody was then added in wash buffer and incubated for 20 min at RT. The samples were kept on ice and, when possible, in the dark. They were washed once with wash buffer and resuspended in PBS (1 ml). Flow cytometry was performed on a BD Accuri C6 (BD Biosciences). Sample data were gated to exclude cell debris, resulting in the detection of a healthy cell population with 10,000 detectable events. Data were analyzed using FlowJo software (www.flowjo.com).

## RESULTS

### Kidney organoids face a developmental plateau in prolonged culture

Kidney organoids were generated from hiPSCs (**Figure 1a**) using an established differentiation protocol [10] with minor adaptations as described by van den Berg and colleagues [19]. A monolayer of hiPSCs was differentiated to renal progenitor cell populations for 7 days. The differentiated cells were subsequently aggregated and transferred to a transwell filter (day 7) to further develop as organoids. The organoids matured over 18 days (day 7+18), as dictated by the protocol, during which renal structures formed. We hypothesized that culture prolongation could be beneficial; however culturing the organoids until day 7+25 resulted in a developmental plateau (**Figure 1a**), in which no further development was apparent. To show that culture prolongation did not negatively affect the presence of renal structures, the organoids were grown until day 7+18 and day 7+25. After this encapsulation, the organoids were characterized for the presence of glomerular structures (Nephrin; NPHS1), proximal tubules (LTL), loop of Henle (NKCC2; SLC12A1), distal tubules (E-cadherin; ECAD) and interstitial cells (Homeobox protein Meis 1/2/3; MEIS1/2/3) (**Figure 1b**). Because culture prolongation showed no loss of renal structures, we sought another explanation for the observed developmental plateau. Therefore, we performed a Masson’s trichrome staining [20, 21]. While kidney organoids at day 7+18 had a homogeneous distribution of glomerular and tubular-like structures, the organoids at day 7+25 showed an expanded interstitial region accompanied by an accumulation of ECM and the manifestation of a stromal cell population (**Figure 1c**).

**Figure 1.**
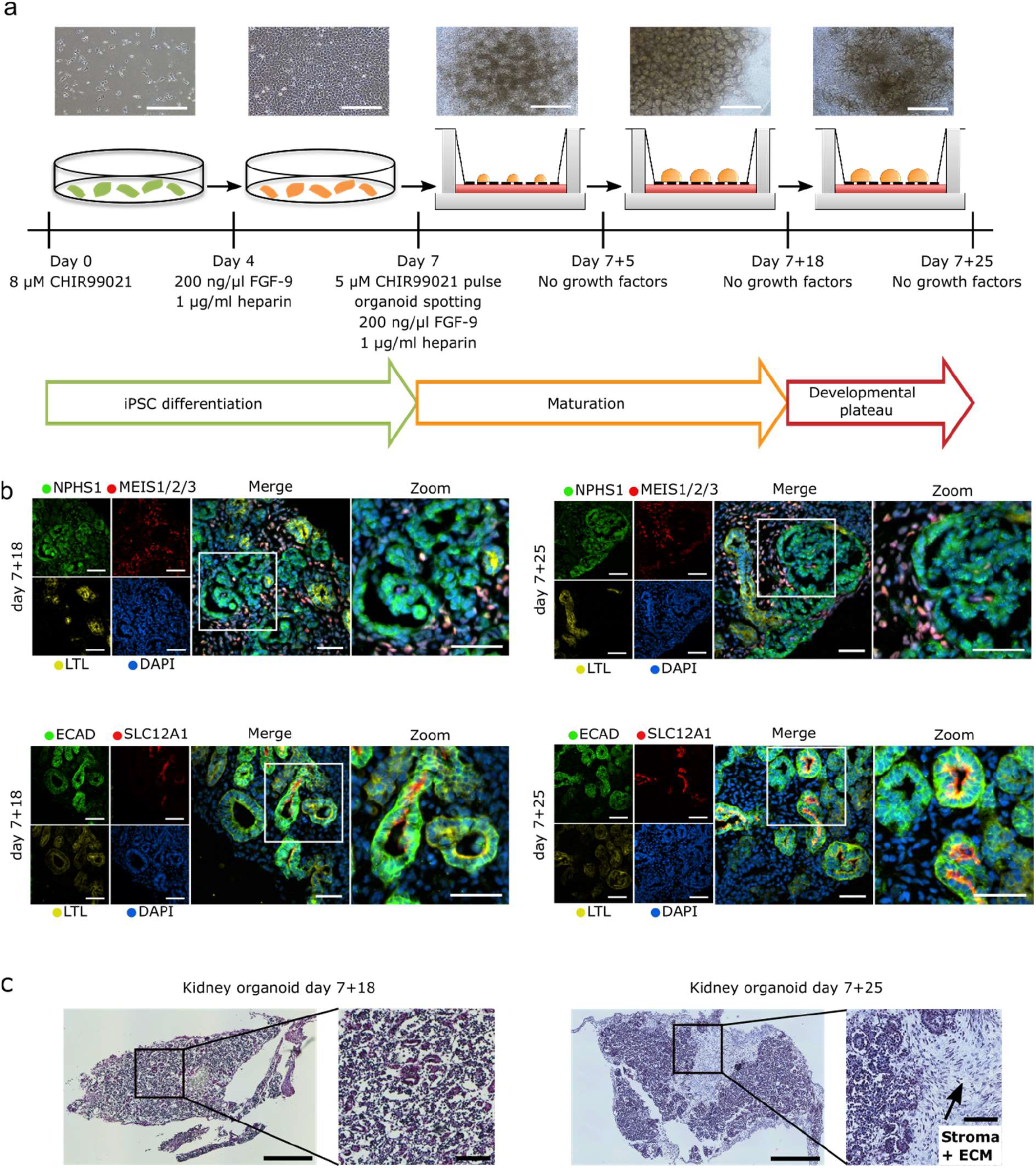
a) Overview of the directed differentiation protocol used to generate kidney organoids. The iPSCs were differentiated for 7 days, after which they were grown and matured as kidney organoids for 18 days. Beyond 18 days, the organoids faced a developmental plateau. Above the schematic, images show the culture at the respective timepoints. Scale bars: 200 μm and 100 μm for first two and last three images, respectively. While organoids aged day 7+25 showed no further maturation, the presence of the different segments of the nephron: glomeruli (Nephrin; NPHS1), proximal tubules (LTL), loop of Henle (NKCC2; SLC12A1), distal tubules (E-cadherin; ECAD) and interstitial cells (Homeobox protein Meis 1/2/3; MEIS1/2/3) was similar to those of organoids at day 7+18. DAPI staining (blue) for nuclei. Scale bars: 50 μm. c) Masson’s trichrome staining of organoids at day 7+18 and day 7+25 showed an expanded interstitial region in the older organoids. This location was accompanied by an accumulation of extracellular matrix and the presence of a stromal cell population. Scale bars: 500 μm for the left images of each set and 100 μm for the zoomed panels.

### Aging organoids in vitro express specific non-renal collagen subtypes

The importance of ECM composition on (stem) cell fate and tissue morphogenesis has already been appreciated [22]. Therefore, we hypothesized that understanding the expression profile of key ECM proteins would help us to understand the developmental plateau that occurred between kidney organoids at day 7+18 and at day 7+25 of culture (**Figure 1a**). Thus, we looked for proteomic changes in the ECM upon culture prolongation (day 7+25) compared to that of day 7+18 cultures. Three biological replicates of human kidney organoids from unique differentiation rounds, aged day 7+18 and day 7+25 were ECM-enriched by depleting cytoplasmic and nuclear proteins (**Figure 2a–b**), then analyzed by tandem mass spectrometry. Extensive data quality control (**Figure S2a–c**) and filtering for valid values yielded a list of 958 proteins identified in at least two of three biological replicates. To analyze for differentially expressed proteins, a two-sample t-test was performed comparing the replicates from day 7+18 with day 7+25 and log-fold changes (LFC) were subsequently calculated (**Figure 2c**). Gene ontology enrichment analysis of differentially expressed proteins (false discovery rate=0.05, LFC=2.0) showed an enrichment of four subnetworks strongly linked to matrix remodeling and collagen expression (**Figure 2d**). More specifically, there was a statistically significant increased expression of collagen types 1, 2 and 6 in the day 7+25 organoids (*e.g.,* COL1A1, abundance ratio=6.850, p-value=1.643E-12; COL2A1, abundance ratio=4.839, p-value=7.309E-09 and COL6A1, abundance ratio=2.799, p-value=1.942E-03) (**Figure 2e, Table 1**), which was confirmed by Western blot (**Figure S2d**).

**Figure 2.**
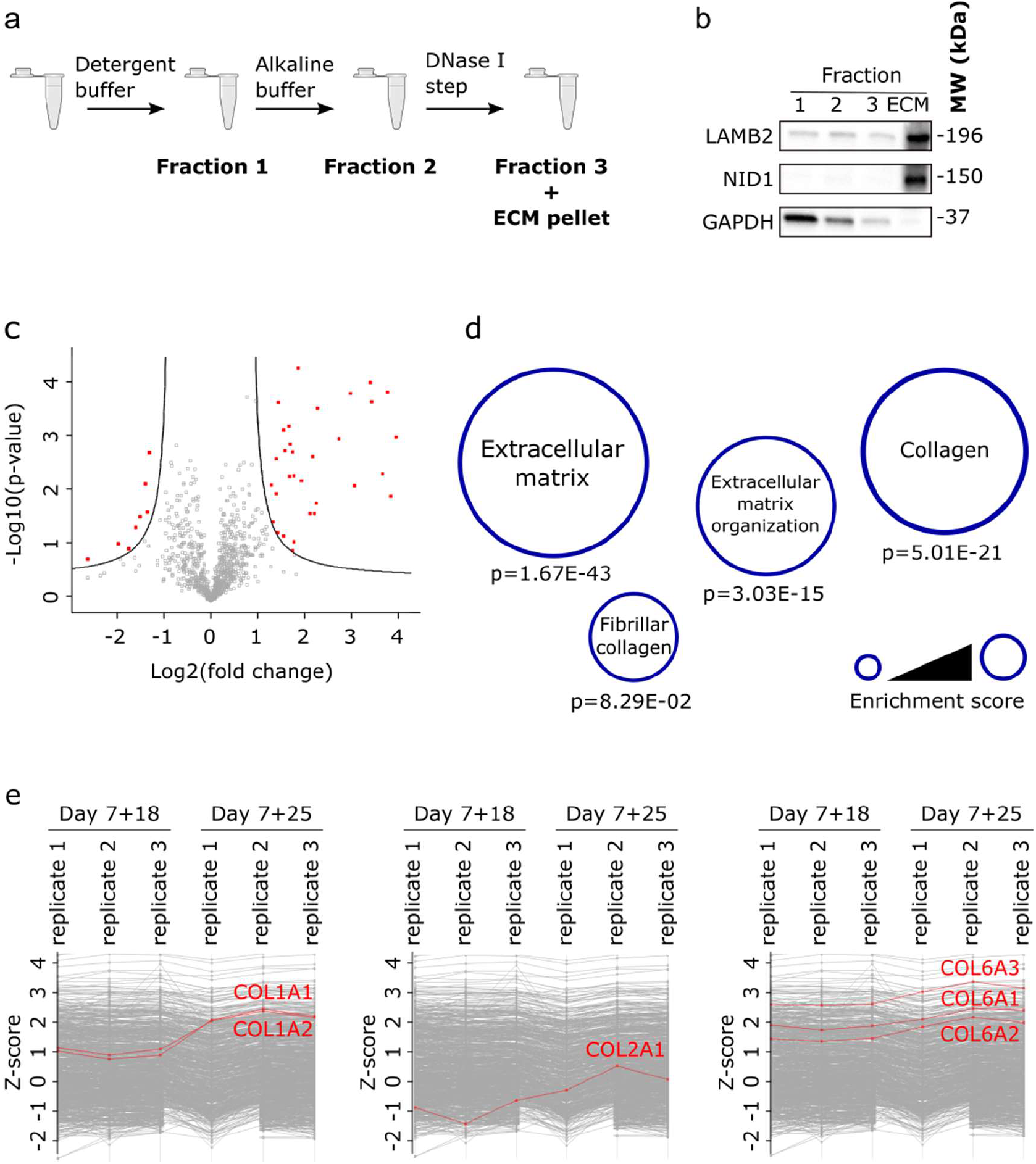
a) Overview of the optimized decellularization protocol to enrich for ECM proteins. b) Western blot validation of the decellularization protocol used for the ECM-enrichment. LAMB2 and NID1 were used as ECM markers and GAPDH as a cytosolic marker. c) Volcano plot of mass spectrometry results showed the differentially expressed proteins (false discovery rate=0.05, log-fold change=2.0). d) Gene ontology analyses showed an enrichment for ECM remodeling and collagen expression. e) Z-score protein expression plots of specific subtypes of type 1, 2 and 6 collagen showed an increase in expression in kidney organoids at day 7+25 compared to those at day 7+18.

**Table 1.** Overview of all ECM and related proteins identified by mass spectrometry analyses of kidney organoids aged day 7+18 and day 7+25. Abundance ratios were calculated to compare protein abundances between the two time points. The p-value was calculated using the Benjamini-Hochberg correction for the false discovery rate.

| **Core matrisome** |  |  |  |  |  |
| --- | --- | --- | --- | --- | --- |
| **Glycoproteins** |  |  | **Collagens** |  |  |
| Gene symbol | Abundance ratio (day 7+25/7+18) | p-value | Gene symbol | Abundance ratio (day 7+25/7+18) | p-value |
| AGRN | 0.615 | 1.707E-01 | COL1A1 | 6.850 | 1.643E-12 |
| DPT | only day 7+25 | 3.353E-16 | COL1A2 | 8.094 | 7.072E-15 |
| EMILIN1 | 1.722 | 3.441E-01 | COL2A1 | 4.839 | 7.309E-09 |
| EMILIN3 | 3.003 | 6.819E-04 | COL4A1 | 0.955 | 9.023E-01 |
| FBN1 | 1.744 | 3.247E-01 | COL4A2 | 0.998 | 9.306E-01 |
| FBN2 | 1.433 | 7.221E-01 | COL5A2 | 2.605 | 2.755E-02 |
| FBN3 | 0.829 | 7.369E-01 | COL5A3 | 1.899 | 3.266E-01 |
| FN1 | 2.555 | 7.699E-03 | COL6A1 | 2.799 | 1.942E-03 |
| LAMA1 | 0.898 | 8.097E-01 | COL6A2 | 2.891 | 1.163E-03 |
| LAMA5 | 0.546 | 5.401E-02 | COL6A3 | 3.107 | 3.771E-04 |
| LAMB1 | 0.595 | 1.262E-01 | COL9A1 | only day 7+25 | 3.353E-16 |
| LAMB2 | 0.588 | 1.134E-01 | COL11A1 | 6.113 | 9.426E-11 |
| LAMC1 | 0.629 | 2.083E-01 | COL12A1 | 4.419 | 2.648E-07 |
| LTBP1 | 1.505 | 6.846E-01 | COL14A1 | 8.184 | 7.072E-15 |
| LTBP4 | 1.223 | 9.187E-01 | COL18A1 | 0.645 | 2.469E-01 |
| MATN4 | 1.987 | 2.721E-01 | COL21A1 | 2.654 | 1.301E-02 |
| MFAP1 | 0.604 | 2.140E-01 | COL26A1 | 1.218 | 9.213E-01 |
| MFAP2 | 1.599 | 4.891E-01 |  |  |  |
| MFAP4 | 3.630 | 1.235E-03 | **Proteoglycans** |  |  |
| NID1 | 0.450 | 4.954E-03 |  |  |  |
| NID2 | 0.685 | 3.334E-01 | ASPN | 20.256 | 3.353E-16 |
| NPNT | 0.708 | 3.833E-01 | BGN | 3.579 | 2.608E-05 |
| POSTN | only day 7+25 | 3.353E-16 | DCN | 13.289 | 3.353E-16 |
| PXDN | 1.175 | 9.359E-01 | FLNA | 1.418 | 7.367E-01 |
| SBSPON | 0.659 | 3.242E-01 | FMOD | 1.739 | 3.266E-01 |
| TGFBI | 1.437 | 7.420E-01 | GPC4 | 1.766 | 2.989E-01 |
| THBS1 | 8.982 | 3.535E-16 | HAPLN1 | 18.854 | 3.353E-16 |
| TNC | 3.496 | 4.001E-05 | LUM | 4.100 | 1.828E-05 |
| VWA1 | 1.523 | 7.100E-01 | OGN | 12.674 | 3.353E-16 |
|  |  |  | PRELP | 1.869 | 2.412E-01 |
|  |  |  | VCAN | 1.785 | 2.890E-01 |
| **Matrisome associated** |  |  |  |  |  |
| **ECM-regulator** |  |  | **Extracellular space** |  |  |
| Gene symbol | Abundance ratio (day 7+25/7+18) | p-value | Gene symbol | Abundance ratio (day 7+25/7+18) | p-value |
| CPXM1 | 1.116 | 9.805E-01 | ANXA1 | 1.779 | 2.890E-01 |
| CSTB | 1.390 | 8.097E-01 | ANXA2 | 1.898 | 1.980E-01 |
| CTSB | 1.255 | 9.043E-01 | ANXA5 | 2.139 | 5.884E-02 |
| TIMP3 | 1.207 | 9.250E-01 | APCS | 2.491 | 1.425E-02 |
|  |  |  | C1QBP | 1.130 | 9.701E-01 |
| **Secreted factors** |  |  | C3 | 1.016 | 9.613E-01 |
|  |  |  | C9 | 1.212 | 9.251E-01 |
| MDK | 1.052 | 9.691E-01 | CLU | 1.701 | 3.483E-01 |
| S100A6 | 1.656 | 4.245E-01 | PPIA | 1.326 | 8.200E-01 |
| S100A10 | 6.553 | 4.361E-09 |  |  |  |
| S100A11 | 1.360 | 7.998E-01 |  |  |  |

An increased expression of specific ECM components, associated with an abnormal local accumulation is typically observed in kidney pathologies [23, 24]. Indeed, we observe a similar phenotype characterized by the increased expression of collagens type 1a1, 6a1 and fibronectin (FN), whose expression is a fibrosis hallmark. More specifically, these fibrotic proteins were found in the interstitial region of aged kidney organoids containing aSMA-positive myofibroblasts (**Figure 3a**), correlating with our previous observations discussed in **Figure 1c**. To understand the kinetics of this collagen deposition in the kidney organoids, we studied the expression profile of type 1a1 and 6a1 collagen over time (days 7+6, +10, +14, +18, +21 and +25) and found a gradual increase as the organoids aged starting from day 7+14 (**Figure 3b and S3**). More specifically, type 1a1 collagen expression was absent before day 7+14 and type 6a1 collagen expression was low. We could confirm these observations by analyzing single cell RNA (scRNA) sequencing datasets from literature [13]. Comparing the distribution of cell populations over time showed a gradual increase for populations expressing the collagen isoforms, together with aSMA and FN (**Figure 3c**). Therefore, we hypothesized that around day 7+14, a (pre-)fibrotic phenotype in kidney organoids emerges.

**Figure 3.**
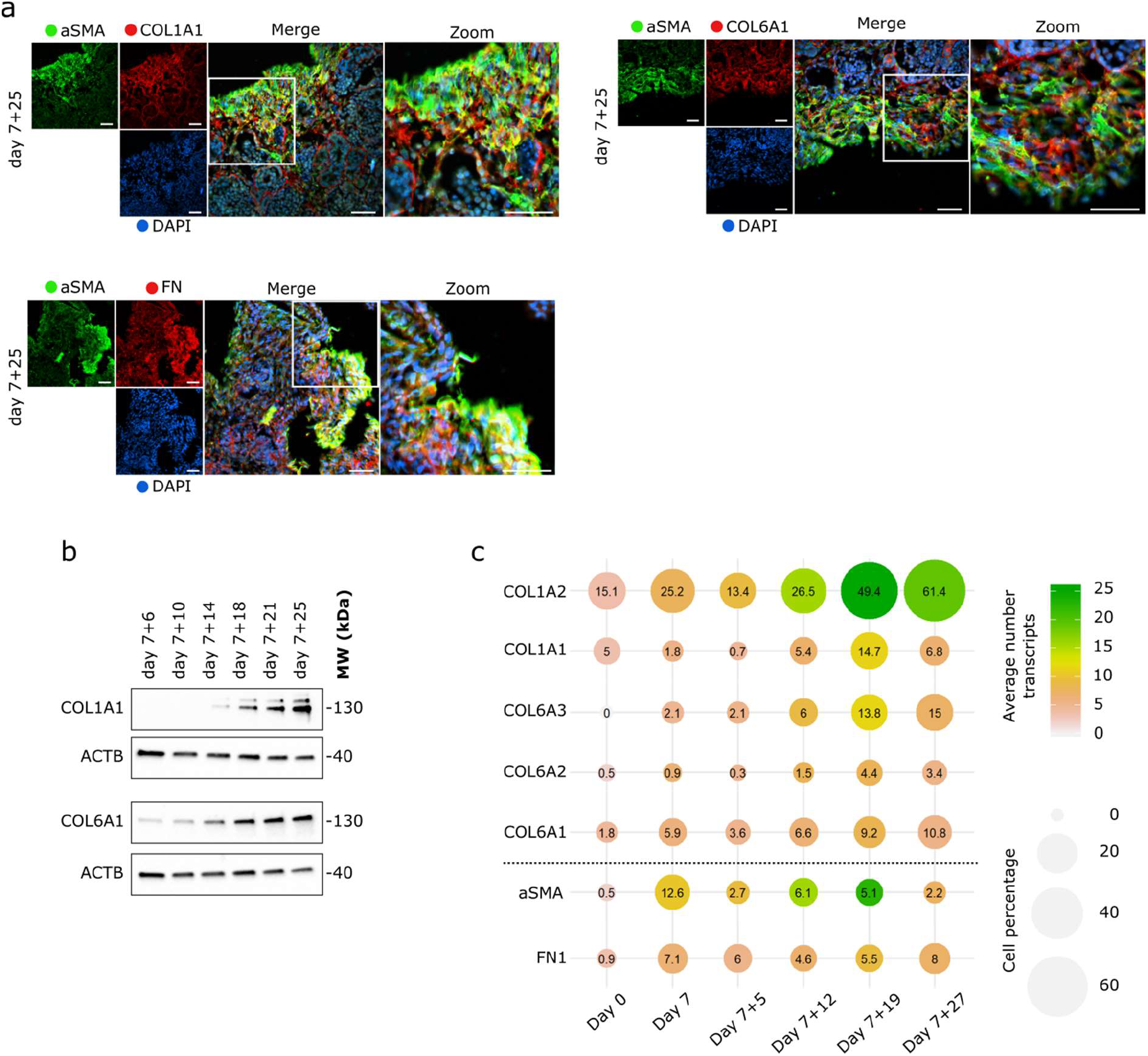
a) Organoids aged day 7+25 showed the presence of a myofibroblast cell population that is responsible for the increased expression of COL1A1, COL6A1 and FN. DAPI staining (blue) for nuclei. Scale bars: 50 μm. b) Both type 1a1 collagen (COL1A1) and type 6 collagen (COL6A1) expression increased in kidney organoids over time. B-actin (ACTB) was used as loading control. Analyzing single cell RNA (scRNA) sequencing datasets from literature [13] showed that cell populations expressing the different COL1 and COL6 isoforms increase when kidney organoids age, as do the fibrosis markers aSMA and fibronectin (FN1).

### A thiol-ene cross-linked alginate was synthesized

When seeking a hydrogel that could modulate the ECM, we began with the observation that the kidney organoids in this study are cultured on an air–liquid interface, which does not represent the natural environment during human kidney development. Therefore, we hypothesized that encapsulating the organoids in a soft hydrogel could reduce the increased expression of collagens type 1a1 and 6 but also the fibrosis markers (FN and aSMA) (**Figure 4a**).

**Figure 4.**
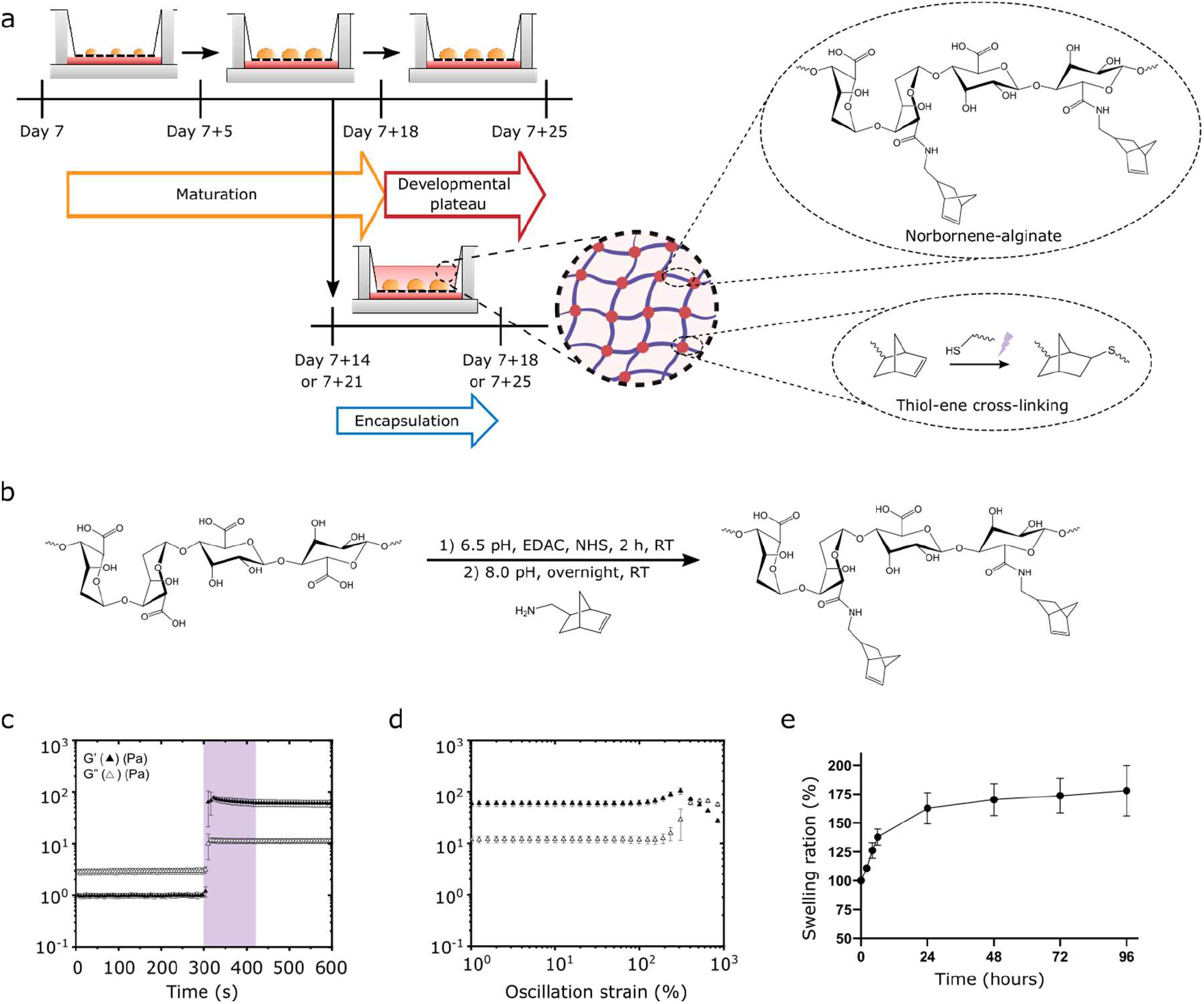
a) Schematic of hydrogel-encapsulated organoid culture. Organoids were encapsulated in a hydrogel at day 7+14 or 7+21 and were cultured for four additional days, until day 7+18 or 7+25, respectively. The hydrogel was composed of norbornene-functionalized alginate crosslinked with a 4-arm thiol via thiol-ene coupling. b) Synthesis schematic of NB-alginate via NHS-ester chemistry. c) *In situ* photo rheometry measurement of the hydrogel (300 s no UV, 120 s with 365 nm UV and 180 s no UV). The hydrogel was formed when exposed to UV light and had a final shear modulus of 60 Pa. G’: storage modulus and G”: loss modulus. d) Strain sweep curve showed slight strain stiffening and a yield point at 250% strain. e) Swelling test of the thiol-ene cross-linked hydrogel shows 175% swelling over a period of 3 days in the simulated culture conditions.

We chose to use alginate, which is a naturally derived, biocompatible, biodegradable, FDA-approved component, and non-adhesive hydrogel [25, 26] that forms an ECM-like environment [26] and allows free diffusion of most nutrients in lower concentration (<2 wt%) hydrogels [27–30]. By covalently modifying the alginate with norbornene (NB-Alg), a UV cross-linkable hydrogel can be created via thiol-ene chemistry. This hydrogel platform allows tunable mechanical properties (without Ca^2+^) and the ability to introduce bioactivity [21]. NB-Alg was synthesized with NHS-chemistry (confirmed by H1-NMR and GPC to contain 3.5% functionalisation, **Figure 4 and S5**). NB-Alg hydrogels were fabricated by cross-linking a 2% (w/v) solution of NB-Alg with a 4-arm PEG thiol with UV light [21] (**Figure 4b**). Rheometry showed an increase of G’ to 10^2^ Pa and G” to 10^1^ Pa after photo exposure and cross-linking (**Figure 4c**), confirming hydrogel formation. Gelation of the hydrogel was observed after 30 sec of UV exposure (purple region in **Figure 4c**), so this time was chosen to encapsulate the organoids. The shear modulus was observed to be constant over a wide frequency regime (**Figure S6a**). Strain sweep measurements showed a slight strain stiffening (of +40 Pa) and a strain at break of approximately 250% (**Figure 4e**). The hydrogels formed a soft (60 Pa shear modulus, ~180 Pa Young’s modulus) and non-adhesive environment into which the organoids could be easily encapsulated.

### Hydrogel encapsulation did not affect organoid viability or morphology

Kidney organoids were encapsulated in NB-Alg hydrogels at day 7+14 and cultured for 4 additional days (**Figure 4a**). Importantly, no structural changes in organoid morphology were observed when comparing brightfield images of encapsulated organoids with organoids cultured on the air–liquid interface at the same time points (**Figure S6e**). The viability of cells at the interface of the organoid and the hydrogel (*e.g.*, the upper surface of the organoid) and within the organoids (*e.g.*, measured from a vertical section) was unchanged whether encapsulated in the NB-Alg hydogel or cultured on the air–liquid interface (**Figure S6d**). This result implies no negative effect of UV exposure or hydrogel encapsulation on the viability of the organoids. The volume of the hydrogel swelled over 4 days to approximately 1.75 times its initial volume (**Figure 4h**), an increase that did not affect the organoids’ structure (**Figure 5a**). In addition, we observed the unchanged presence of physiologically relevant cell populations and structures of the organoids when encapsulated, which we based on the following markers: (Nephrin; NPHS1), proximal tubules (LTL), loop of Henle (NKCC2; SLC12A1), distal tubules (E-cadherin; ECAD) and interstitial cells (Homeobox protein Meis 1/2/3; MEIS1/2/3) (**Figure 5a**).

**Figure 5.**
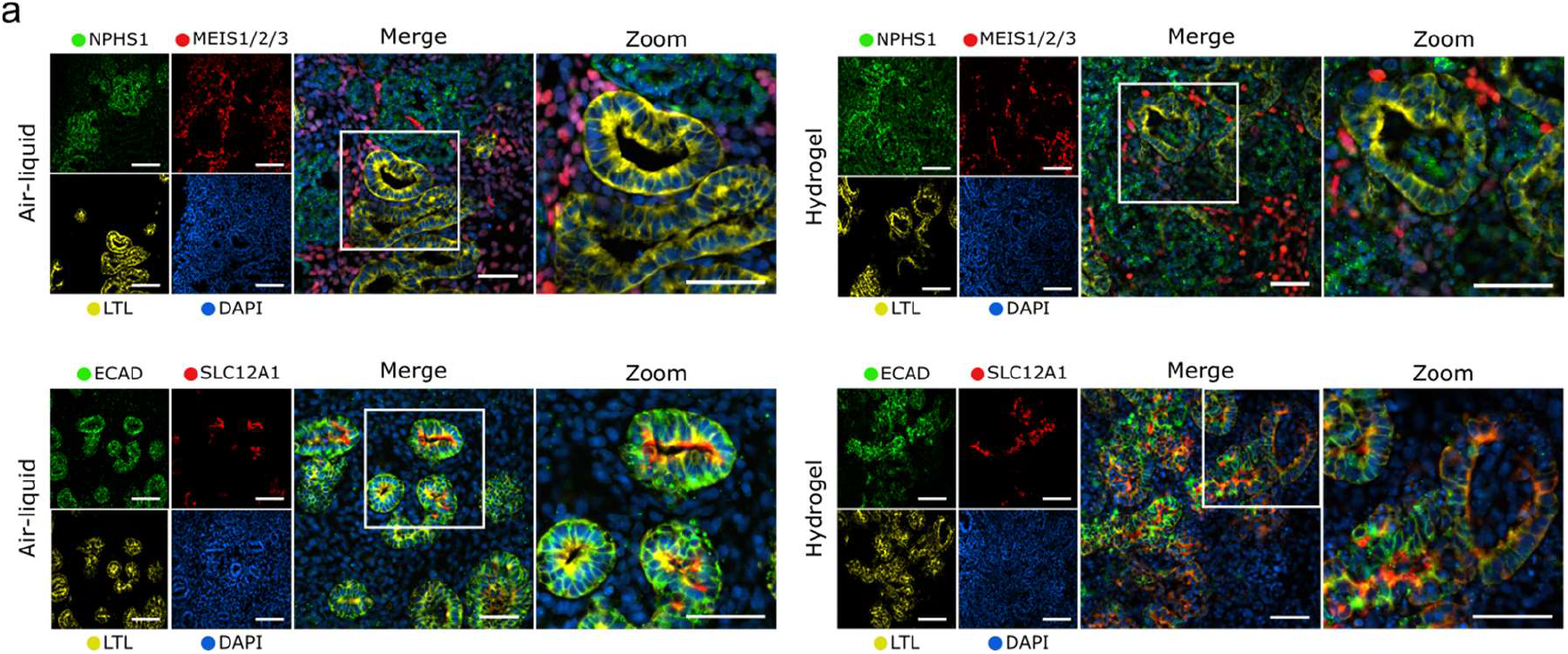
a) Encapsulated organoids until day 7+18 (right image panel) did not show differences in the presence of the different segments of the nephron: glomeruli (Nephrin; NPHS1), proximal tubules (LTL), loop of Henle (NKCC2; SLC12A1), distal tubules (E-cadherin; ECAD) and interstitial cells (Homeobox protein Meis 1/2/3; MEIS1/2/3) when compared to organoids grown on an air–liquid interface until day 7+18 (left image panel). DAPI staining (blue) for nuclei. Scale bars: 50 μm.

### Kidney organoid encapsulation reduces type 1a1 collagen deposition

To determine whether hydrogel encapsulation effectively reduced collagen expression (and thus could slow the progression of a fibrotic-like phenotype), we recovered the organoids from the hydrogel at day 7+18 of culture (**Figure 6a**) before enriching for the ECM proteins as described (**Figure 3a**). The expression of type 1a1 collagen was diminished in the encapsulated organoids compared to those cultured on the air–liquid interface (**Figure 6b, S8**). We found no consistent change in the expression of type 6a1 collagen (**Figure 6b, S8**). These results indicate a selective modifying effect of the hydrogel encapsulation on type 1a1 collagen deposition.

**Figure 6.**
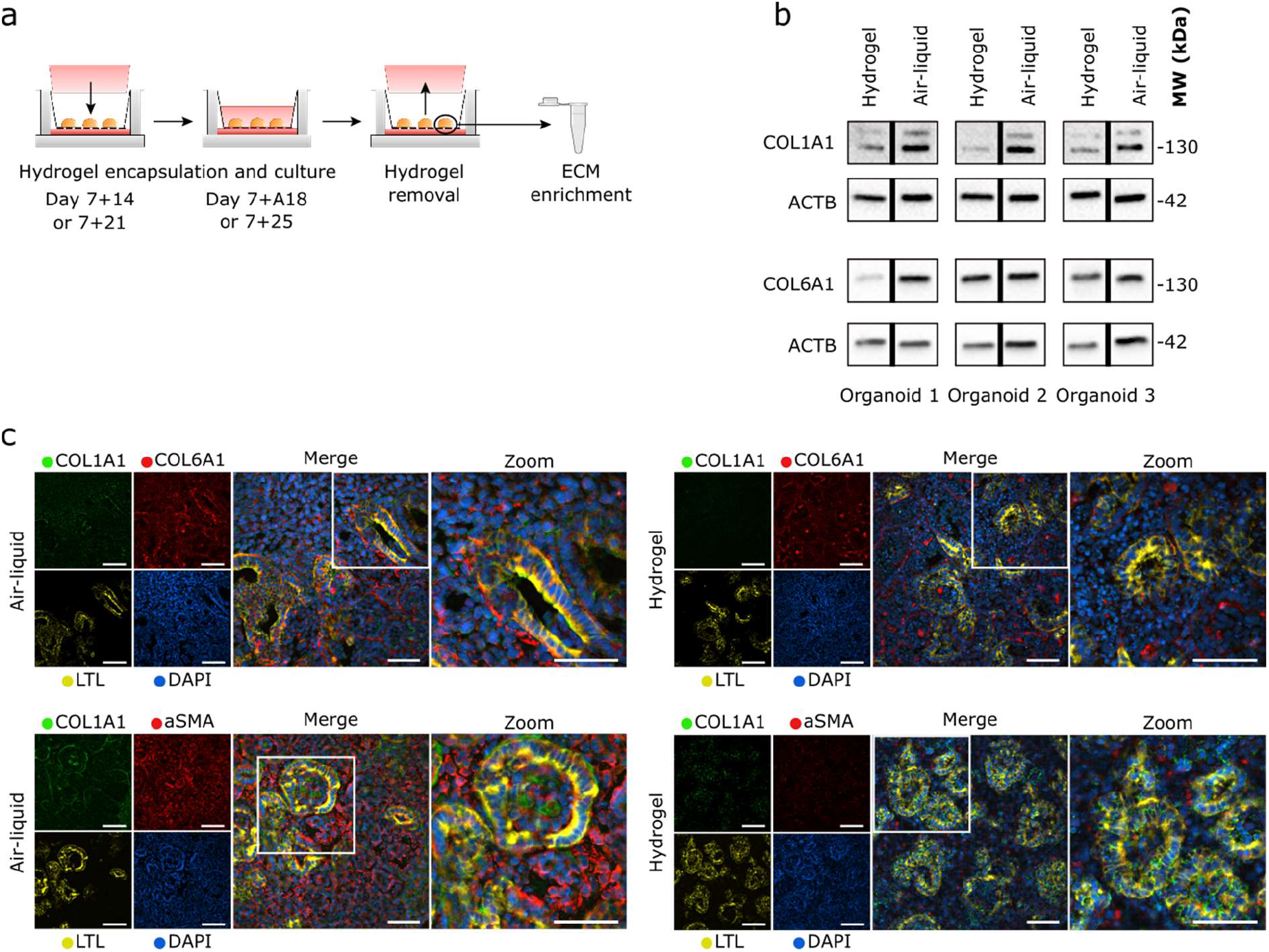
a) Schematic of organoid retrieval from the hydrogel after encapsulation. b) Western blotting of ECM proteins type 1a1 collagen and type 6a1 collagen in organoids encapsulated in the hydrogel from day 7+14 to 7+18 compared to those from the air–liquid interface culture at day 7+18. A reduction of type 1a1 collagen was observed from all three sampled organoids encapsulated in the hydrogels. B-actin (ACTB) was used as a loading control. c) Immunohistochemistry for collagen type 1a1 and 6a1, αSMA and LTL on horizontally sectioned organoids encapsulated from day 7+14 to 7+18 compared to organoids cultured at the air–liquid interface at day 7+18. Reduced expression of type 1a1 collagen and αSMA was observed in the hydrogel-encapsulated organoids. Scale bars: 50 μm.

In order to find the most efficient time for encapsulation to reduce the abnormal ECM deposition, we investigated the encapsulation effect on organoids at a later culture time point. More specifically, the kidney organoids were encapsulated at day 7+21 and cultured until day 7+25 (**Figure 6a**). Again, a reduction in type 1a1 collagen was observed in the encapsulated organoids compared to those cultured on the air–liquid interface at day 7+25 (**Figure S7a**). This reduction was similar to what we observed before in organoids encapsulated at day 7+18 (**Figure 6b**), indicating that the time point of encapsulation, and therefore the existing amount of collagen type 1a1 at the time of encapsulation, may not affect the ability to reduce the increased type 1a1 collagen deposition in kidney organoids.

To exclude the possibility that type 1a1 collagen was removed with the hydrogel upon analysis, sections of the organoids encapsulated in the hydrogel were stained for type 1a1 collagen. The images showed decreased type 1a1 collagen expression within the encapsulated organoids compared to those cultured on the air–liquid interface, comparable to the reduction observed on the Western blots. Interestingly, type 6a1 collagen depostition remains unchanged when comparing air-liquid interface and encapsulated organoids. Areas of dense type 1a1 collagen expression were observed in organoids cultured on the air– liquid interface, while a more diffuse and overall lower amount was observed when the organoids were encapsulated in the hydrogel (**Figure 6c, S7b**). Interestingly, the expression of the fibrotic marker (aSMA) was also reduced, similarly to type 1a1 collagen, in the encapsulated organoids (**Figure 6c, S7b**).

## DISCUSSION

Being able to create kidney organoids derived from patient or donor hiPSCs holds great promise as a novel source of functional nephrons for patients suffering from end-stage renal diseases. Despite recent breakthroughs, these organoids reveal a fibrotic phenotype preventing them from being translated to the clinic. In this study, we investigated if *in vitro* culture prolongation combined with a hydrogel encapsulation could decrease the collagen and αSMA expression to prolong the culture of the organoids without the emergence of the fibrotic phenotype. Using a proteomic approach we found that prolonged *in vitro* culture conditions of kidney organoids resulted in the increased expression of three specific collagen subtypes (type 1a1, 2 and 6a1) and the fibrotic marker αSMA. Organoid encapsulation in a soft, thiol-ene cross-linked alginate hydrogel decreased the expression of these subtypes, thereby pointing to a possible effect of the biomaterial on reducing the fibrotic phenotype in aging kidney organoids.

Creating a replacement therapy for patients with CKD requires functional organoids that should not be expressing the hallmarks of renal fibrosis. We observe an increased deposition of specific types of collagens such as type 1a1, 2 and 6, which has been reported to be characteristic of fibrillary-like scar tissue in renal fibrosis and CKD [23, 24, 31, 32]. In a healthy kidney, the expression of these types is very low, or even absent, but upon kidney damage or disease, activated fibroblasts turn into myofibroblasts and (over)express specific types of collagens to deposit them in the renal interstitium [15, 16]. An excessive accumulation of ECM components is a hallmark of fibrotic lesions (as shown by the abnormally expanded interstitial region in older organoids, **Figure 2a**) and leads to a life-threatening pathological condition by compromising the organ’s function [17]. Being able to improve *in vitro* organoid maturation without the appearance of fibrosis is crucial to fully exploit them as promising clinical solutions. Tissue engineers have developed numerous approaches to influence cell fate and morphology by engineering the microenvironment. Hydrogels are of high interest due to their similarities with the ECM architecture and their high water content (>90%). Many studies have looked at the impact of hydrogel encapsulation on different types of organoids [33]; such as the intestine [34–37], pancreas [38, 39], neural tube [40] and liver [38]. However, the use of hydrogels is still limited in the kidney organoid field [41, 42].

For the time period of encapsulation we tested (4 days), reductions were consistent for collagen type 1a1 but not for collagen type 6. This specificity has also been reported in methacrylated alginate hydrogels where softer gels (0.7 kPa) also reduced the type 1 collagen expression and stiffer gels (2.5 kPa) had a positive effect on isotropic collagen organization, with no effect on type 3 collagen [43]. Soft porcine kidney decellurized ECM hydrogels of stiffnesses between 0.02 and 0.1 kPa have been previously reported to support human glomerular endotheial cell viability and gene expression [44]. Furthermore, the addition of type 1 collagen to increase the stiffness of human kidney cortex decellularized ECM hydrogels (0.25 kPa) has been reported to lead to an active cell state when encapsulating human kidney peritubular microvascular endothelial cells [45]. Besides hydrogel stiffness, the relative relaxation time has been observed to affect type 1 collagen deposition. For example, the encapsulation of chondrocytes in hydrazone covalently adapatible network PEG hydrogels resulted in lower levels of type 1 collagen in hydrogels with faster relaxation times, with no effect on type 2 collagen [43, 46]. Interestingly, tuning the hydrogel’s mechanical properties (specifically stiffness or relaxation time) did not affect the expression and modulation of type 2 [46] or type 3 collagen [43] in these studies, which is similar to our observations for type 6 collagen. Finally, the reduction of type 1 collagen could also be caused by increased collagen degradation activity by matrix metalloproteinases, as has been reported for a cartilage-specific degradable poly(ethylene) glycol hydrogel used for encapsulating chondrocytes [47].

The alginate hydrogels in this study contained no cellular anchors, such as RGD, to interact with the cells. These nonadhesive properties therefore preclude adhesive mechanotransduction events. However, both our study and the two studies discussed previously, point to a possible effect of the nonadhesive encapsulation on extracellular protein expression [43, 46]. This postulates an indirect effect of the surrounding environment’s stiffness and relaxation time on cellular behaviour. The hydrogels we report here have a shear modulus of 60 Pa (corresponding Young’s modulus of 180 Pa); this stiffness is significantly different from the human kidney, which has a Young’s modulus ranging from 5 to 10 kPa [48]. Instead, the hydrogel envrionment employed in our study more closely mimics the perinephric adipose tissue formed around the renal capsule, which has a Young’s modulus of approximately 0.5 kPa [48–50]. This raises the question of whether we may have overlooked the organ’s surrounding environment *in vivo* when reconstructing organs *in vitro*. Supporting this idea, kidneys have been found to be enveloped with brown adipose tissue during development [51]. This fat encapsulation provides the developing kidney with a soft extracellular surrounding, possibly regulating complex events during nephrogenesis. Interestingly, recent studies pointed to the positive effect on the maturation status of kidney organoids by conducting transplantation experiments under the mouse renal capsule or onto the chick chorioallantoic membrane [19, 42]. Therefore, future research could be focussed on the design of biomaterials with similar properties to the human renal capsule to unravel the importance of dynamic interactions, relaxation time, stiffness and fibrous morphology properties.

## CONCLUSION

Differentiated kidney organoids from induced pluripotent stem cells hold promise as a treatment for patients with kidney diseases. Before these organoids can be translated to the clinic, shortcomings regarding their extracellular composition need to be overcome. We showed that simple encapsulation into a soft biomaterial could control the organoids’ ECM, potentially diminishing or delaying the fibrotic trajectory that organoids undergo during aging in vitro. This step forward gives organoids the chance to mature and makes them more suitable for clinical applications.

## ACKNOWLEDGMENTS

This work is supported by the partners of Regenerative Medicine Crossing Borders (RegMed XB), a public–private partnership that uses regenerative medicine strategies to cure common chronic diseases. This collaboration is financed by the Dutch Ministry of Economic Affairs by means of the PPP Allowance made available by the Top Sector Life Sciences & Health to stimulate public–private partnerships. MBB and VLSL also thank the Dutch Province of Limburg, FLCM and MBB acknowledge funding from NWO under project agreement 731.016.202 “DynAM”, and TR and VLSL acknowledge funding from the European Research Council (ERC) under the European Union’s Horizon 2020 research and innovation programme (grant agreement No 694801). The authors thank Christian Freund (hiPSC core facility, LUMC, Leiden, the Netherlands) for providing the hiPSC line (LUMC0072iCTRL01), Nathalie Groen for performing the scRNA sequencing analyses and the Maastricht MultiModal Molecular Imaging Institute–Division of Imaging Mass Spectrometry for performing the mass spectrometry.

## DATA AVAILABILITY

The raw and processed data required to reproduce these findings are available to download from https://dataverse.nl/privateurl.xhtml?token=04ca9a0b-3fe4-4bab-838d-04d53167cbda.

## SUPPORTING FIGURES

**Figure S1.**
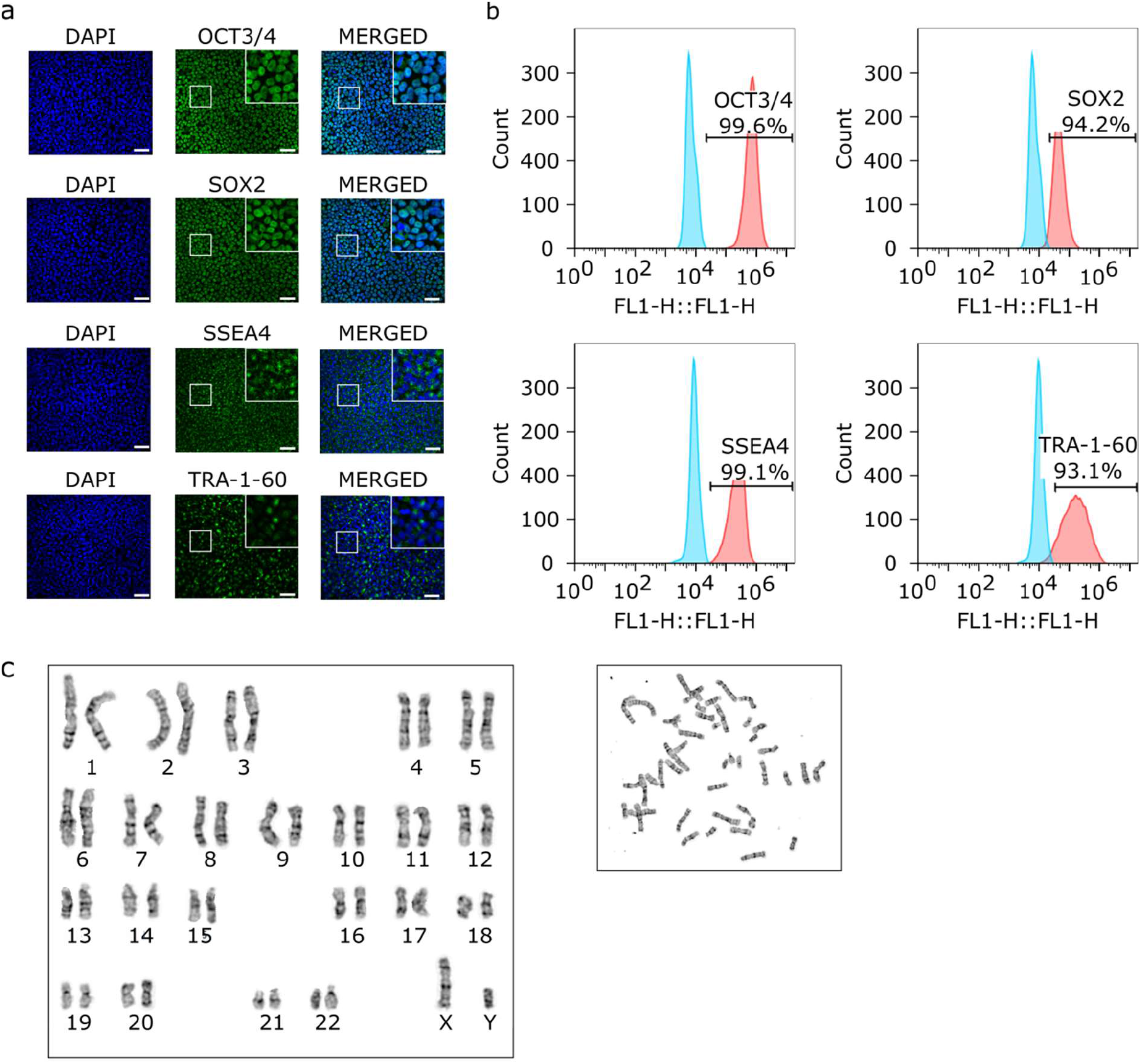
a) Immunocytochemistry shows the pluripotency status of the iPSC cell line used to generate kidney organoids by the high expression of the pluripotency markers (middle column): OCT3/4, SOX2, SSEA4 and TRA-1-60. For each image, the inset in the upper right corner shows the respective boxed area at 40× magnification. DAPI staining (left column) of nuclei. Right column shows the merged images. Scale bars: 50 μm. b) Flow cytometry analyses showed a high percentage of cells expressing the pluripotency markers. c) Karyogram of the studied iPSC cell line showed a normal karyotype.

**Figure S2.**
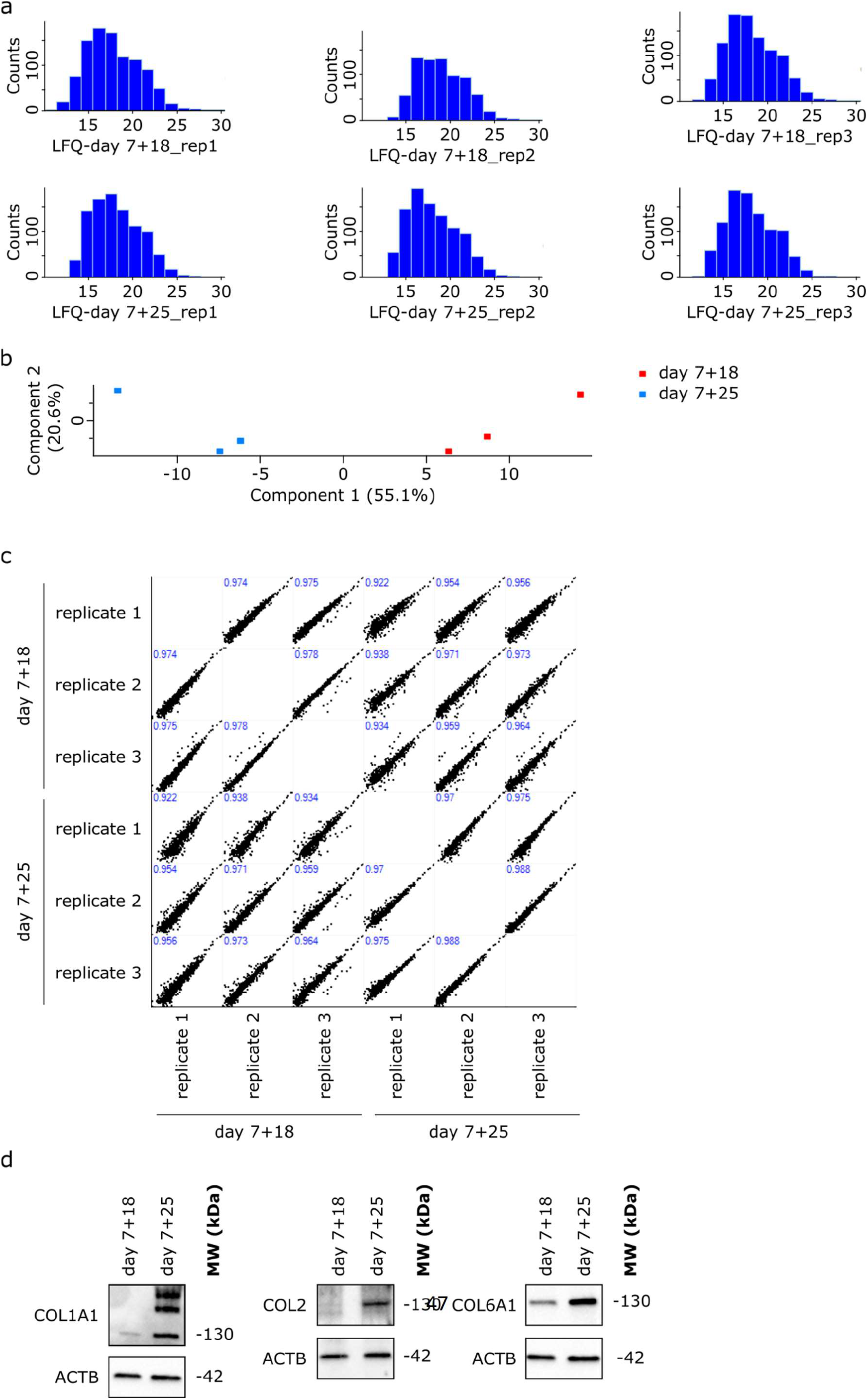
a) Histograms showing a normal distribution for the data obtained by mass spectrometry. b) Principal component analysis showed the clustering of the day 7+18 replicates that is distinct from the day 7+25 replicates. c) Multi scatter plots were used to assess for the correlation between samples. Pearson’s correlation coefficient (show in blue) is high between all samples and between the replicate groups. d) Validation of the obtained mass spectrometry data by Western blotting for a small set of identified differentially expressed proteins: COL1A1, COL2 and COL6A1. B-actin (ACTB) was used as a loading control.

**Figure S3.**
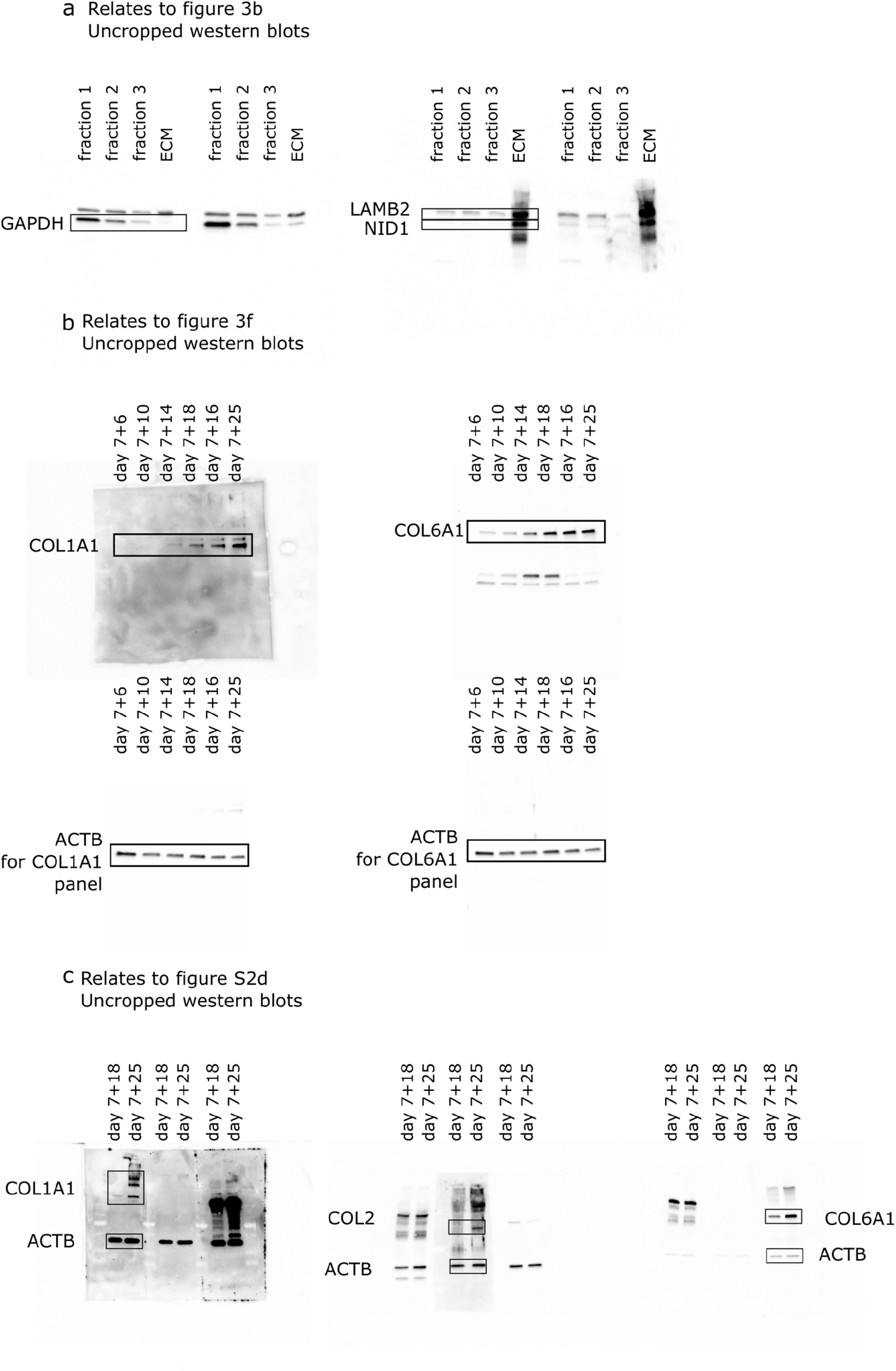
Uncropped Western blot images for corresponding figures as labeled.

**Figure S4.**
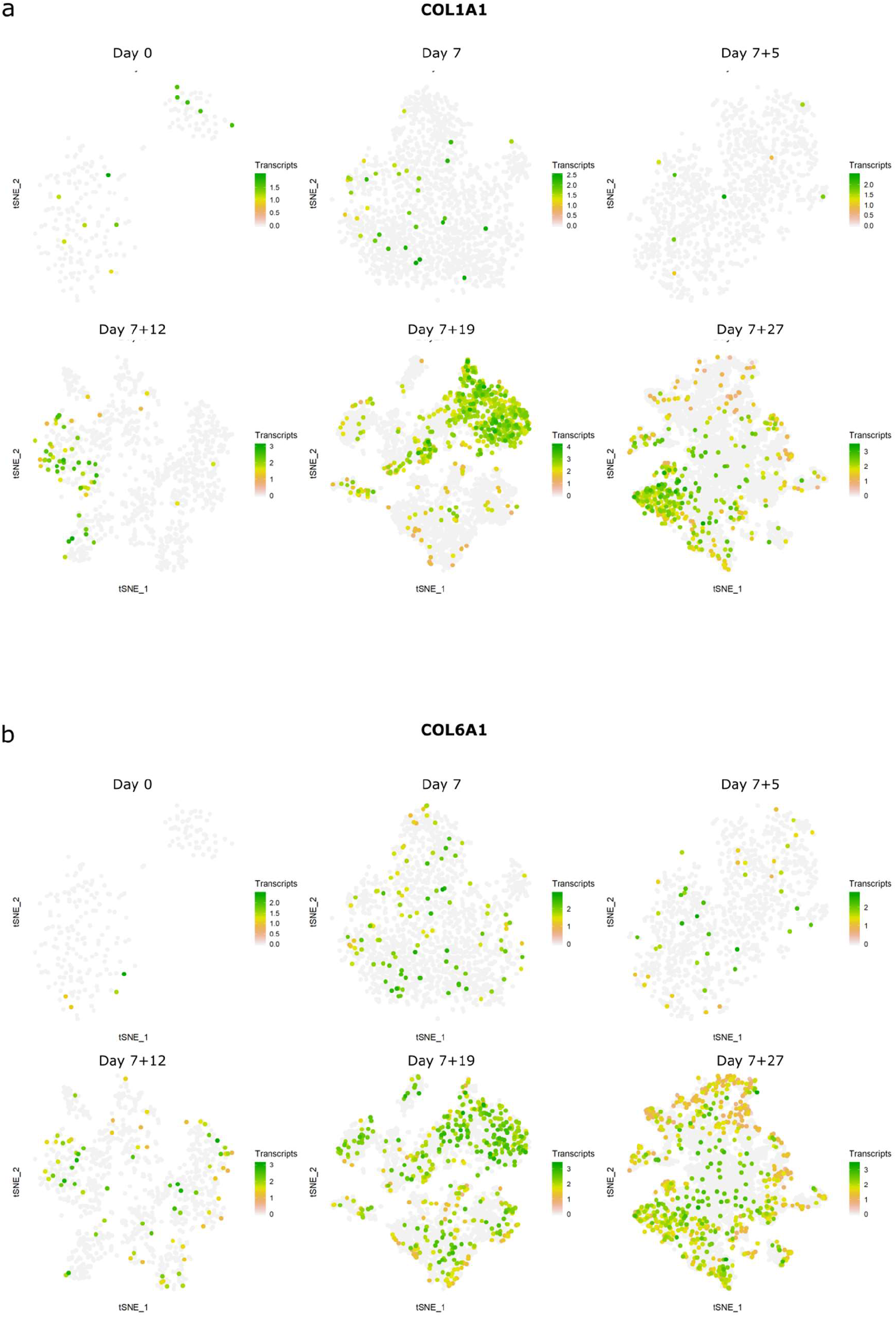
Normalized gene expression of markers of interest are shown in tSNE space. Cell populations expressing (a) COL1A1 and (b) COL6A1 increase in number when kidney organoid culture was prolonged.

**Figure S5.**
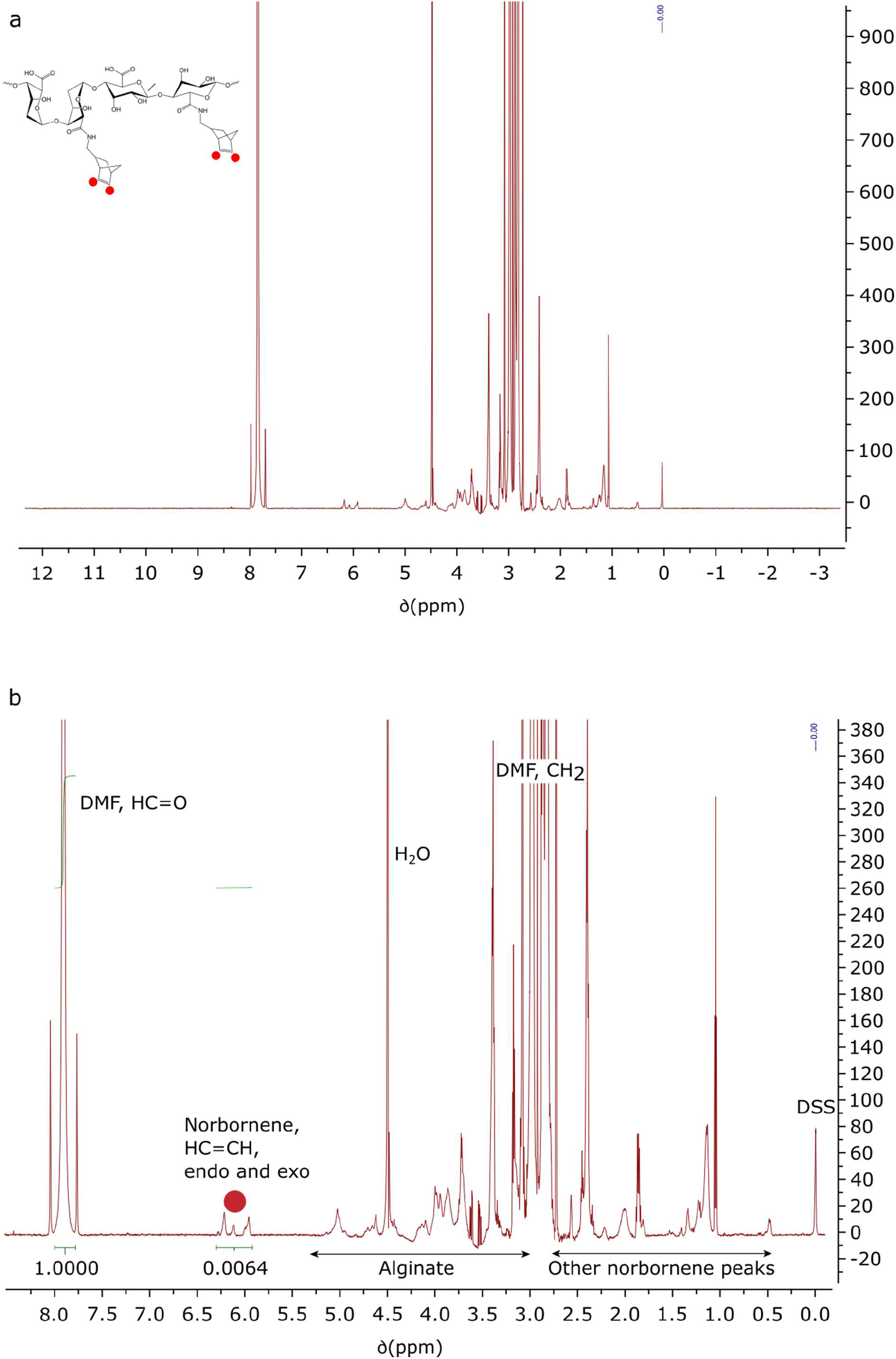
a) Full ^1^H-NMR spectra of norbornene-functionalized alginate. b) Zoom of the ^1^H-NMR spectra of norbornene-functionalized alginate. CH=CH peaks of the norbornene (endo and exo) at 5.95, 6.15, 6.25 and 6.27 ppm, in D_2_O. DMF: internal standard (0.2 M). For calculations of functionalization, see Table S1.

**Figure S6.**
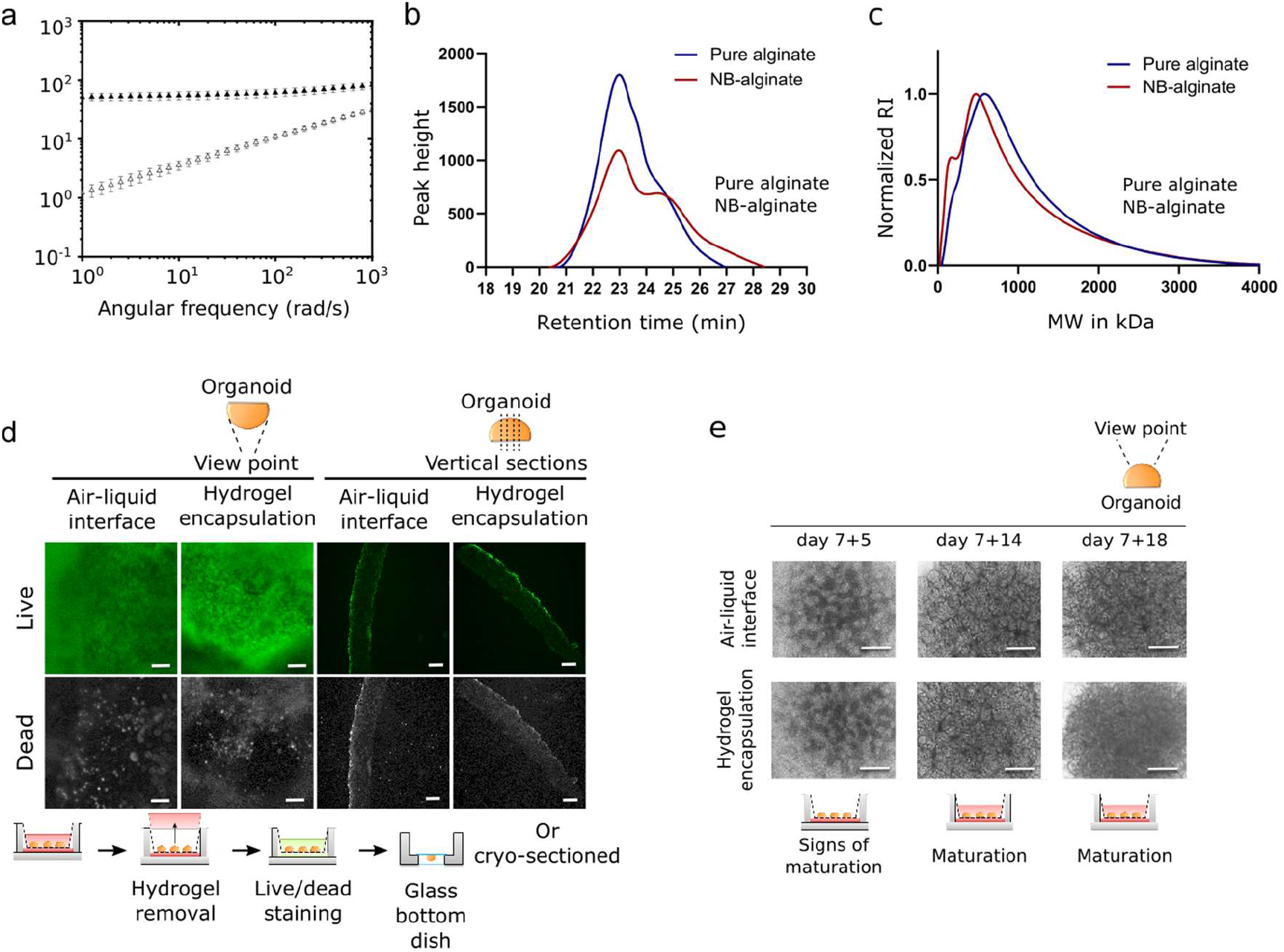
a) Frequency sweep curves at 1% strain showed consistent G’ over a wide time range. b–c) GPC plots of NB-alginate vs. pure alginate for retention time vs. peak height (f) and MW vs. normalized RI, Mn, Mw, etc. values in Table S2 (c). d) Live/dead staining (EthD1/Calcein AM) fluorescence imaging of whole mounted organoids (two left columns) and vertical cryosections (two right columns). N=4 for each condition. e) Brightfield images of organoid structures at day 7+5 showing the first signs of nephrogenesis, day 7+14 before encapsulation, and day 7+18 after 3 days of encapsulation. N=4, scale bars: 100 μm. Scale bars: 100 μm.

**Figure S7.**
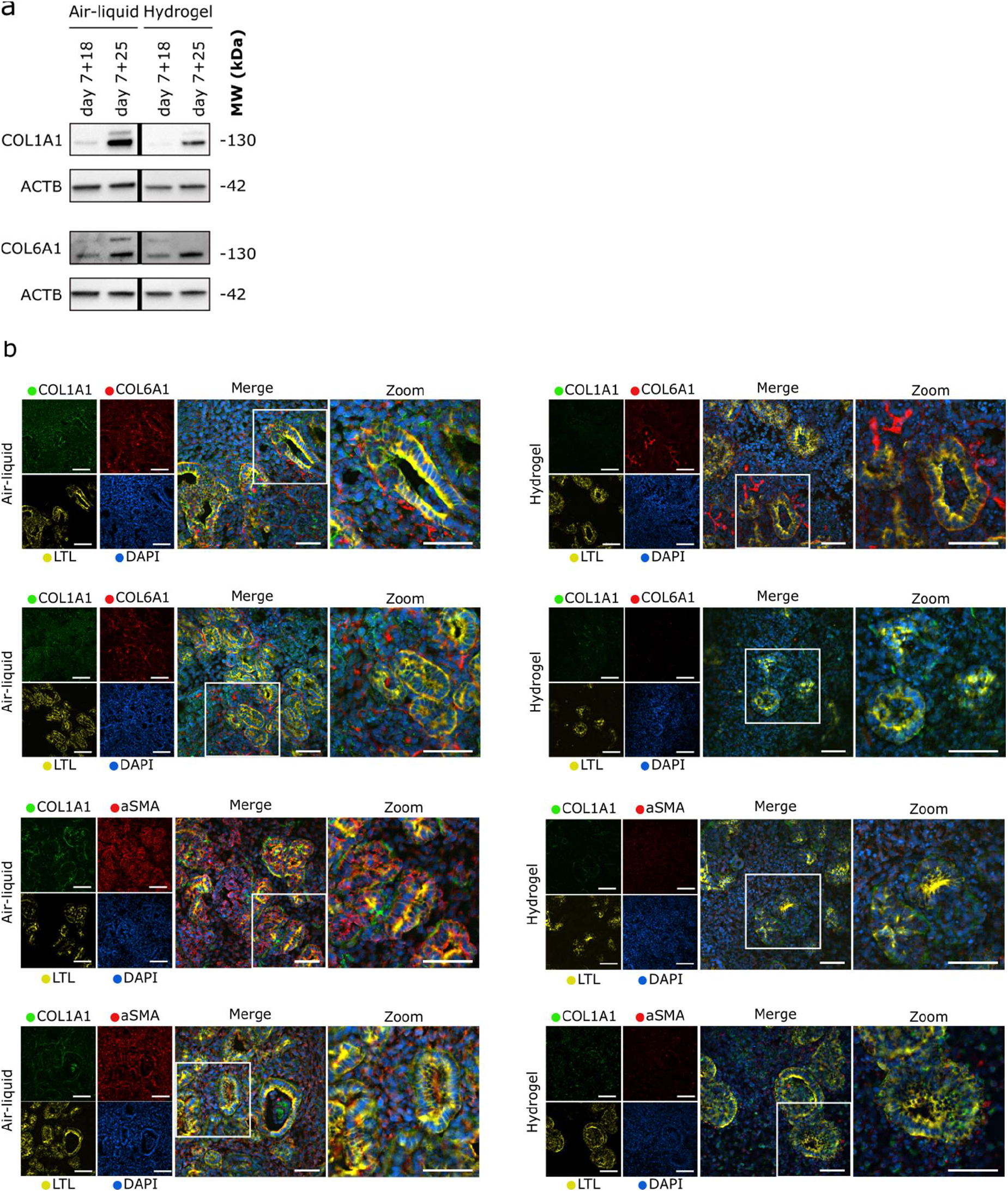
a) Western blotting of ECM proteins type 1a1 and type 6 collagen from organoids encapsulated in the hydrogel from day 7+14 to 7+18 and from day 7+21 to 7+25 of culture compared to air-liquid interface at the same respective timepoints. B-actin (ACTB) was used as a loading control. b) Immunohistochemistry for collagen type 1a1 and 6a1, aSMA and LTL in horizontally sectioned organoids from two unique differentiation rounds, encapsulated from day 7+14 to 7+18 compared to organoids cultured at the air–liquid interface at day 7+18. Scale bars: 50 μm.

**Figure S8.**
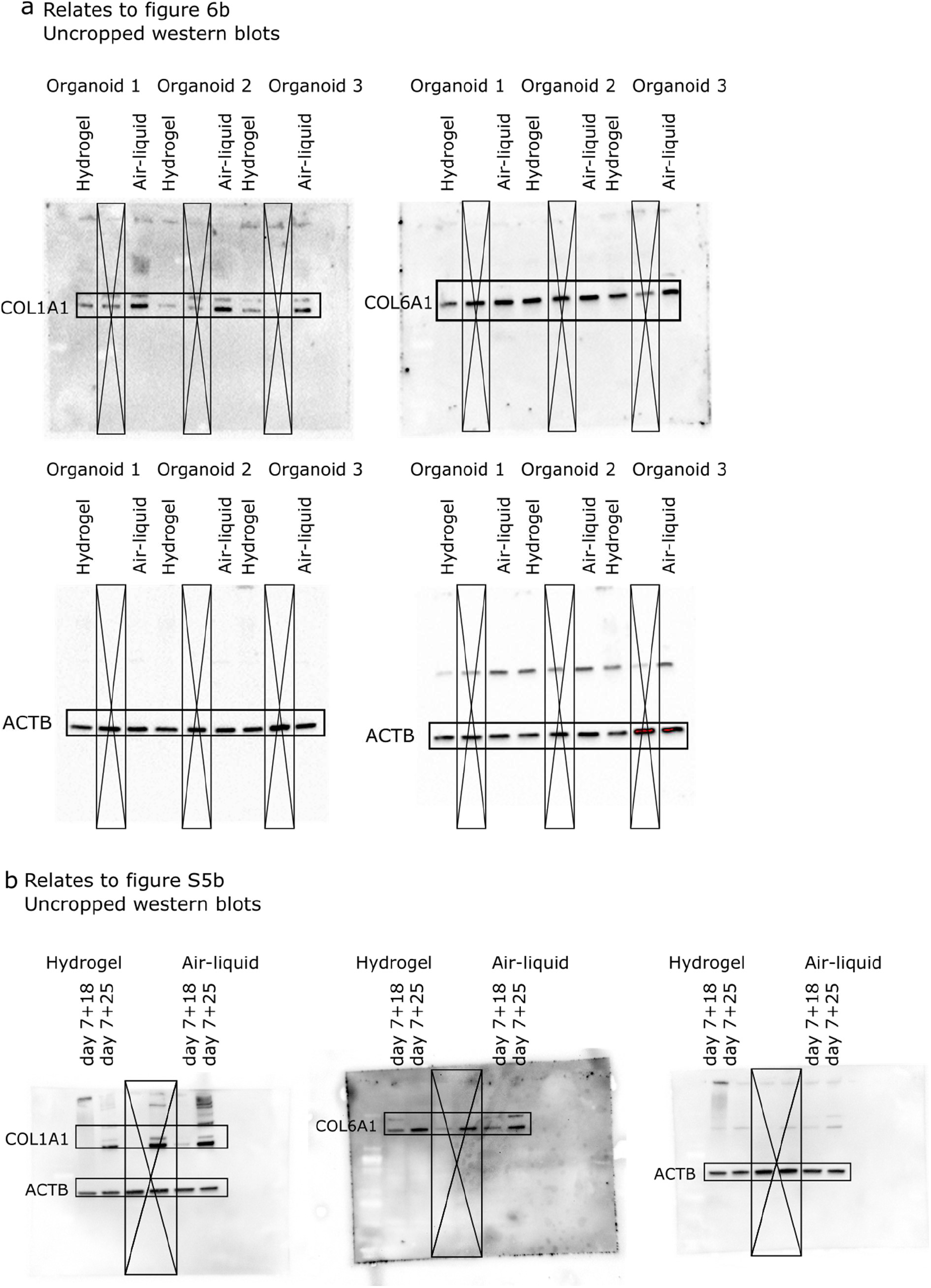
Uncropped Western blot images used for corresponding figures as labeled.

**Table S1.**
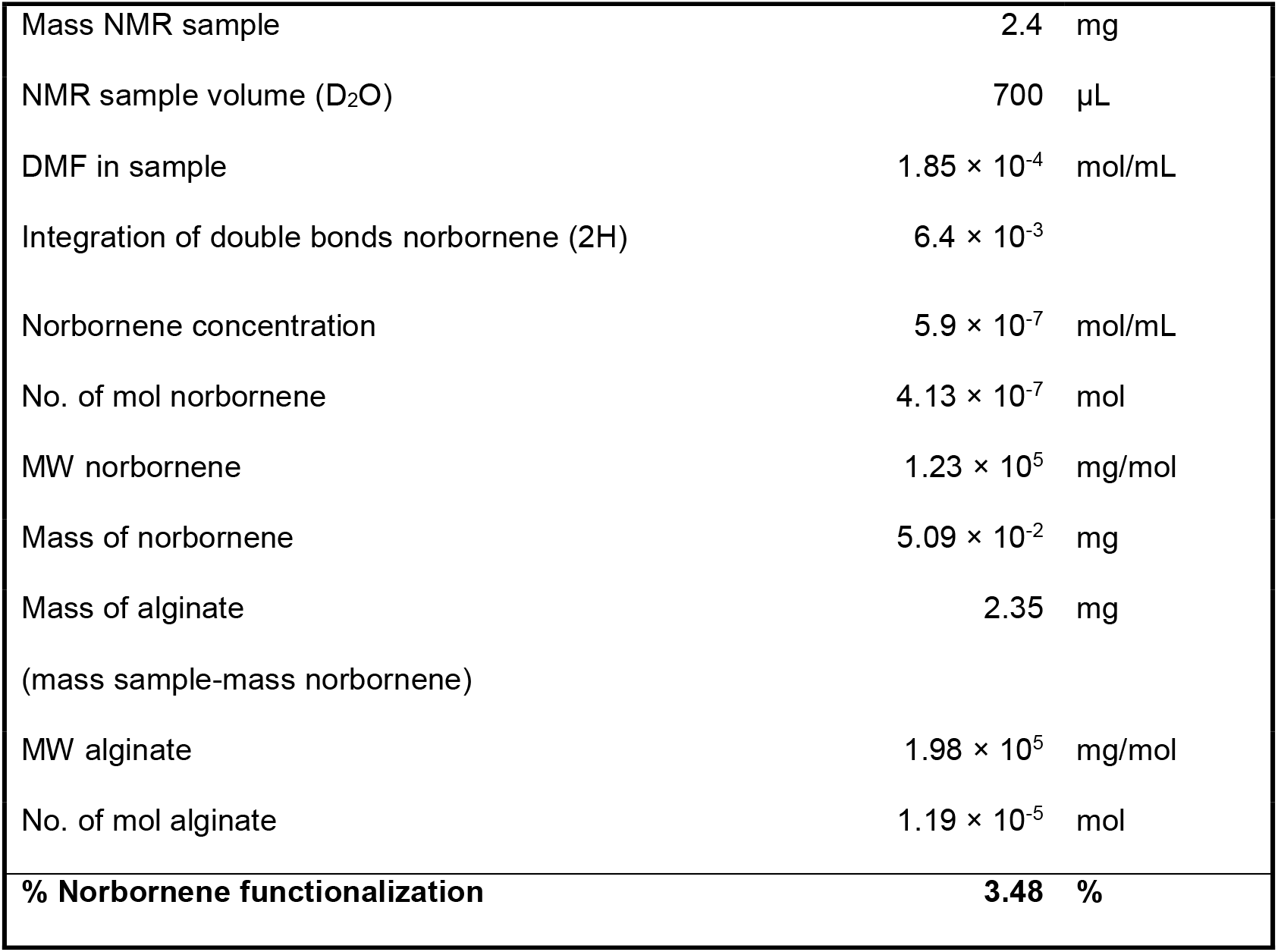
Calculations for estimation of degree of norbornene functionalization on the alginate with ^1^H-NMR integrals using DMF as an internal standard.

|  |  |  |
| --- | --- | --- |
| Mass NMR sample | 2.4 | mg |
| NMR sample volume (D<sub>2</sub>O) | 700 | μL |
| DMF in sample | $1.85 \times 10^{-4}$ | mol/mL |
| Integration of double bonds norbornene (2H) | $6.4 \times 10^{-3}$ | |
| Norbornene concentration | $5.9 \times 10^{-7}$ | mol/mL |
| No. of mol norbornene | $4.13 \times 10^{-7}$ | mol |
| MW norbornene | $1.23 \times 10^5$ | mg/mol |
| Mass of norbornene | $5.09 \times 10^{-2}$ | mg |
| Mass of alginate | 2.35 | mg |
| (mass sample-mass norbornene) |  |  |
| MW alginate | $1.98 \times 10^5$ | mg/mol |
| No. of mol alginate | $1.19 \times 10^{-5}$ | mol |
| **% Norbornene functionalization** | **3.48** | **%** |

**Table S2.** Molecular weight averages by GPC. Number average (M_n_), Weight average (M_w_), and dispersity (M_w_/M_n_) for pure alginate and norbornene-functionalized (NB-) alginate.

| | $M_n$ [kDa] | $M_w$ [kDa] | $M_w/M_n$ [Đ] |
| --- | --- | --- | --- |
| **Pure Alg** | $2.6 \times 10^5$ | $4.6 \times 10^5$ | 1.8 |
| **NB-Alg** | $1.7 \times 10^5$ | $4.4 \times 10^5$ | 2.6 |

## REFERENCES

[1] N.R. Hill, S.T. Fatoba, J.L. Oke, J.A. Hirst, C.A. O’Callaghan, D.S. Lasserson, F.D.R. Hobbs, Global prevalence of chronic kidney disease – a systematic review and meta-analysis, PLOS One 11(7) (2016) e0158765.

[2] P. Romagnani, G. Remuzzi, R. Glassock, A. Levin, K.J. Jager, M. Tonelli, Z. Massy, C. Wanner, H.J. Anders, Chronic kidney disease, Nat. Rev. Dis. Primers 3 (2017) 17088.

[3] P. Chang, J. Gill, J. Dong, C. Rose, H. Yan, D. Landsberg, E.H. Cole, J.S. Gill, Living donor age and kidney allograft half-life: implications for living donor paired exchange programs, Clin. J. Am. Soc. Nephrol. 7(5) (2012) 835–841.

[4] M.H. Little, P. Kairath, Regenerative medicine in kidney disease, Kidney Int. 90(2) (2016) 289–299.

[5] F. Schutgens, M.B. Rookmaaker, T. Margaritis, A. Rios, C. Ammerlaan, J. Jansen, L. Gijzen, M. Vormann, A. Vonk, M. Viveen, F.Y. Yengej, S. Derakhshan, K.M. de Winter-de Groot, B. Artegiani, R. van Boxtel, E. Cuppen, A.P.A. Hendrickx, M.M. van den Heuvel-Eibrink, E. Heitzer, H. Lanz, J. Beekman, J.L. Murk, R. Masereeuw, F. Holstege, J. Drost, M.C. Verhaar, H. Clevers, Tubuloids derived from human adult kidney and urine for personalized disease modeling, Nat. Biotechnol. 37(3) (2019) 303–313.

[6] A. Fatehullah, S.H. Tan, N. Barker, Organoids as an in vitro model of human development and disease, Nat. Cell Biol. 18(3) (2016) 246–254.

[7] A. Taguchi, Y. Kaku, T. Ohmori, S. Sharmin, M. Ogawa, H. Sasaki, R. Nishinakamura, Redefining the in vivo origin of metanephric nephron progenitors enables generation of complex kidney structures from pluripotent stem cells, Cell Stem Cell 14(1) (2014) 53–67.

[8] B.S. Freedman, C.R. Brooks, A.Q. Lam, H. Fu, R. Morizane, V. Agrawal, A.F. Saad, M.K. Li, M.R. Hughes, R.V. Werff, D.T. Peters, J. Lu, A. Baccei, A.M. Siedlecki, M.T. Valerius, K. Musunuru, K.M. McNagny, T.I. Steinman, J. Zhou, P.H. Lerou, J.V. Bonventre, Modelling kidney disease with CRISPR-mutant kidney organoids derived from human pluripotent epiblast spheroids, Nat. Commun. 6(1) (2015) 8715–8728.

[9] R. Morizane, A.Q. Lam, B.S. Freedman, S. Kishi, M.T. Valerius, J.V. Bonventre, Nephron organoids derived from human pluripotent stem cells model kidney development and injury, Nat. Biotechnol. 33(11) (2015) 1193–1200.

[10] M. Takasato, P.X. Er, H.S. Chiu, B. Maier, G.J. Baillie, C. Ferguson, R.G. Parton, E.J. Wolvetang, M.S. Roost, S.M. Chuva de Sousa Lopes, M.H. Little, Kidney organoids from human iPS cells contain multiple lineages and model human nephrogenesis, Nature 526(7574) (2015) 564–568.

[11] A. Taguchi, R. Nishinakamura, Higher-order kidney organogenesis from pluripotent stem cells, Cell Stem Cell 21(6) (2017) 730–746.e6.

[12] T. Geuens, C.A. van Blitterswijk, V.L.S. LaPointe, Overcoming kidney organoid challenges for regenerative medicine, npj Regenerative Medicine 5(1) (2020) 8–14.

[13] H. Wu, K. Uchimura, E.L. Donnelly, Y. Kirita, S.A. Morris, B.D. Humphreys, Comparative analysis and refinement of human PSC-derived kidney organoid differentiation with single-cell transcriptomics, Cell Stem Cell 23(6) (2018) 869–881.

[14] M. Takasato, P.X. Er, H.S. Chiu, M.H. Little, Generation of kidney organoids from human pluripotent stem cells, Nat. Protoc. 11(9) (2016) 1681–1692.

[15] S. Meran, R. Steadman, Fibroblasts and myofibroblasts in renal fibrosis, Int. J. Exp. Pathol. 92(3) (2011) 158–167.

[16] R.D. Bulow, P. Boor, Extracellular matrix in kidney fibrosis: more than just a scaffold, J. Histochem. Cytochem. 67(9) (2019) 643–661.

[17] A. Fernández-Colino, L. Iop, M.S. Ventura Ferreira, P. Mela, Fibrosis in tissue engineering and regenerative medicine: treat or trigger?, Adv. Drug Deliv. Rev. 146 (2019) 17–36.

[18] S. Tyanova, T. Temu, P. Sinitcyn, A. Carlson, M.Y. Hein, T. Geiger, M. Mann, J. Cox, The Perseus computational platform for comprehensive analysis of (prote)omics data, Nat. Methods 13(9) (2016) 731–740.

[19] C.W. van den Berg, L. Ritsma, M.C. Avramut, L.E. Wiersma, B.M. van den Berg, D.G. Leuning, E. Lievers, M. Koning, J.M. Vanslambrouck, A.J. Koster, S.E. Howden, M. Takasato, M.H. Little, T.J. Rabelink, Renal subcapsular transplantation of PSC-derived kidney organoids induces neo-vasculogenesis and significant glomerular and tubular maturation in vivo, Stem Cell Rep. 10(3) (2018) 751–765.

[20] A.B.A. Farris C.E., What is the best way to measure renal fibrosis?: A pathologist’s perspective, Kidney Int. Suppl. 4(1) (2014) 9–15.

[21] H.W. Ooi, C. Mota, A.T. ten Cate, A. Calore, L. Moroni, M.B. Baker, Thiol–ene alginate hydrogels as versatile bioinks for bioprinting, Biomacromolecules 19(8) (2018) 3390–3400.

[22] C.M. Nelson, M.M. Vanduijn, J.L. Inman, D.A. Fletcher, M.J. Bissell, Tissue geometry determines sites of mammary branching morphogenesis in organotypic cultures, Science 314(5797) (2006) 298–300.

[23] F. Genovese, A.A. Manresa, D.J. Leeming, M.A. Karsdal, P. Boor, The extracellular matrix in the kidney: a source of novel non-invasive biomarkers of kidney fibrosis?, Fibrogenesis Tissue Repair 7(1) (2014) 4–4.

[24] H. Menn-Josephy, C.S. Lee, A. Nolin, M. Christov, D.V. Rybin, J.M. Weinberg, J. Henderson, R. Bonegio, A. Havasi, Renal interstitial fibrosis: an imperfect predictor of kidney disease progression in some patient cohorts, Am. J. Nephrol. 44(4) (2016) 289–299.

[25] T. Andersen, B.L. Strand, K. Formo, E. Alsberg, B.E. Christensen, Chapter 9 Alginates as biomaterials in tissue engineering, Carbohydrate Chemistry, The Royal Society of Chemistry, Published, 2012, p. 227–258.

[26] J. Sun, H. Tan, Alginate-based biomaterials for regenerative medicine applications, Materials 6(4) (2013) 1285–1309.

[27] W.R. Gombotz, S.F. Wee, Protein release from alginate matrices, Adv. Drug Deliv. Rev. 64 (2012) 194–205.

[28] A. Khavari, M. Nydén, D.A. Weitz, A.J. Ehrlicher, Composite alginate gels for tunable cellular microenvironment mechanics, Sci. Rep. 6 (2016) 30854–30864.

[29] A. Espona-Noguera, J. Ciriza, A. Cañibano-Hernández, L. Fernandez, I. Ochoa, L. Saenz del Burgo, J.L. Pedraz, Tunable injectable alginate-based hydrogel for cell therapy in type 1 diabetes mellitus, Int. J. Biol. Macromol. 107 (2018) 1261–1269.

[30] M.H. Hettiaratchi, A. Schudel, T. Rouse, A.J. García, S.N. Thomas, R.E. Guldberg, T.C. McDevitt, A rapid method for determining protein diffusion through hydrogels for regenerative medicine applications, APL Bioeng. 2(2) (2018) 026110.

[31] A.A. Eddy, Overview of the cellular and molecular basis of kidney fibrosis, Kidney Int. Suppl. 4(1) (2014) 2–8.

[32] S. Buchtler, A. Grill, S. Hofmarksrichter, P. Stockert, G. Schiechl-Brachner, M. Rodriguez Gomez, S. Neumayer, K. Schmidbauer, Y. Talke, B.M. Klinkhammer, P. Boor, A. Medvinsky, K. Renner, H. Castrop, M. Mack, Cellular origin and functional relevance of collagen I production in the kidney, J. Am. Soc. Nephrol. 29(7) (2018) 1859–1873.

[33] M.J. Kratochvil, A.J. Seymour, T.L. Li, S.P. Paşca, C.J. Kuo, S.C. Heilshorn, Engineered materials for organoid systems, Nature Reviews Materials 4(9) (2019) 606–622.

[34] R.L. DiMarco, R.E. Dewi, G. Bernal, C. Kuo, S.C. Heilshorn, Protein-engineered scaffolds for in vitro 3D culture of primary adult intestinal organoids, Biomaterials Science 3(10) (2015) 1376–1385.

[35] N. Gjorevski, M.P. Lutolf, Synthesis and characterization of well-defined hydrogel matrices and their application to intestinal stem cell and organoid culture, Nature Protoc. 12(11) (2017) 2263–2274.

[36] R. Cruz-Acuña, M. Quirós, A.E. Farkas, P.H. Dedhia, S. Huang, D. Siuda, V. García-Hernández, A.J. Miller, J.R. Spence, A. Nusrat, A.J. García, Synthetic Hydrogels for Human Intestinal Organoid Generation and Colonic Wound Repair, Nat. Cell Biol. 19(11) (2017) 1326–1335.

[37] M.M. Capeling, M. Czerwinski, S. Huang, Y.-H. Tsai, A. Wu, M.S. Nagy, B. Juliar, N. Sundaram, Y. Song, W.M. Han, S. Takayama, E. Alsberg, A.J. Garcia, M. Helmrath, A.J. Putnam, J.R. Spence, Nonadhesive alginate hydrogels support growth of pluripotent stem cell-derived intestinal organoids, Stem Cell Rep. 12(2) (2019) 381–394.

[38] L. Broutier, A. Andersson-Rolf, C.J. Hindley, S.F. Boj, H. Clevers, B.-K. Koo, M. Huch, Culture and establishment of self-renewing human and mouse adult liver and pancreas 3D organoids and their genetic manipulation, Nat. Protoc. 11 (2016) 1724–1743.

[39] J. Candiello, T.S.P. Grandhi, S.K. Goh, V. Vaidya, M. Lemmon-Kishi, K.R. Eliato, R. Ros, P.N. Kumta, K. Rege, I. Banerjee, 3D heterogeneous islet organoid generation from human embryonic stem cells using a novel engineered hydrogel platform, Biomaterials 177 (2018) 27–39.

[40] A. Ranga, M. Girgin, A. Meinhardt, D. Eberle, M. Caiazzo, E.M. Tanaka, M.P. Lutolf, Neural tube morphogenesis in synthetic 3D microenvironments, PNAS 113(44) (2016) E6831–E6839.

[41] N. Broguiere, L. Isenmann, C. Hirt, T. Ringel, S. Placzek, E. Cavalli, F. Ringnalda, L. Villiger, R. Züllig, R. Lehmann, G. Rogler, M.H. Heim, J. Schüler, M. Zenobi-Wong, G. Schwank, Growth of epithelial organoids in a defined hydrogel, Adv. Mater. 30(43) (2018) 1801621.

[42] E. Garreta, P. Prado, C. Tarantino, R. Oria, L. Fanlo, E. Martí, D. Zalvidea, X. Trepat, P. Roca-Cusachs, A. Gavaldà-Navarro, L. Cozzuto, J.M. Campistol, J.C. Izpisúa Belmonte, C. Hurtado del Pozo, N. Montserrat, Fine tuning the extracellular environment accelerates the derivation of kidney organoids from human pluripotent stem cells, Nat. Mater. 18(4) (2019) 397–405.

[43] A. Boddupalli, K.M. Bratlie, Collagen organization deposited by fibroblasts encapsulated in pH responsive methacrylated alginate hydrogels, J. Biomed. Mat. Res. Part A 106(11) (2018) 2934–2943.

[44] J. Su, S.C. Satchell, R.N. Shah, J.A. Wertheim, Kidney decellularized extracellular matrix hydrogels: Rheological characterization and human glomerular endothelial cell response to encapsulation, J. Biomed. Mat. Res. Part A 106(9) (2018) 2448–2462.

[45] R.J. Nagao, J. Xu, P. Luo, J. Xue, Y. Wang, S. Kotha, W. Zeng, X. Fu, J. Himmelfarb, Y. Zheng, Decellularized human kidney cortex hydrogels enhance kidney microvascular endothelial cell maturation and quiescence, Tissue Eng. Part A 22(19-20) (2016) 1140–1150.

[46] B.M. Richardson, D.G. Wilcox, M.A. Randolph, K.S. Anseth, Hydrazone covalent adaptable networks modulate extracellular matrix deposition for cartilage tissue engineering, Acta Biomater. 83 (2019) 71–82.

[47] S.C. Skaalure, S. Chu, S.J. Bryant, An enzyme-sensitive PEG hydrogel based on aggrecan catabolism for cartilage tissue engineering, Adv. Healthc. Mater. 4(3) (2015) 420–431.

[48] A.M. Handorf, Y. Zhou, M.A. Halanski, W.-J. Li, Tissue stiffness dictates development, homeostasis, and disease progression, Organogenesis 11(1) (2015) 1–15.

[49] M. Geerligs, G.W. Peters, P.A. Ackermans, C.W. Oomens, F.P. Baaijens, Linear viscoelastic behavior of subcutaneous adipose tissue, Biorheology 45(6) (2008) 677–688.

[50] T.R. Cox, J.T. Erler, Remodeling and homeostasis of the extracellular matrix: implications for fibrotic diseases and cancer, Dis. Models Mech. 4(2) (2011) 165–178.

[51] H. Sacks, M.E. Symonds, Anatomical locations of human brown adipose tissue: functional relevance and implications in obesity and type 2 diabetes, Diabetes 62(6) (2013) 1783–1790.

